# ATP synthase deficiency promotes prolonged hydrogen production in Chlamydomonas

**DOI:** 10.64898/2026.10.08.757524

**Authors:** Jade Rachel Marcus, Tamar Elman, Felix Buchert, Iftach Yacoby

## Abstract

Photosynthetic electron transport is tightly regulated to balance ATP and NADPH production and prevent over-reduction of the photosynthetic machinery. How this balance is maintained under severe ATP limitation, particularly under anoxia, is not fully resolved. Here, we used several deletion mutants of chloroplast ATP synthase subunits from *Chlamydomonas reinhardtii* to examine how electron flow is reorganized under anoxic conditions through the interplay between linear electron flow, cyclic electron flow, and alternative electron sinks. Using a combination of optical spectroscopy methods and membrane inlet mass spectrometry, we characterized changes in photosynthetic electron transport, proton motive force relaxation and gas exchange. Under oxic conditions, the ΔatpH mutant (lacking the c-subunit) showed strong acceptor-side limitation at photosystem I (PSI), whereas *atpC1-2* (lacking the γ-subunit) resembled the wild type. Under anoxia, this limitation was partially relieved, coinciding with sustained electron transport and prolonged hydrogen production. In particular, the Δ*atpH* mutant exhibited increased cyclic electron flow under oxic conditions and maintained PSI activity under anoxia despite impaired ATP synthesis. Prolonged hydrogen production was also observed in *atpC1-2*, which did not share the increased respiration of ΔatpH. Hydrogenase activity under anoxia is consistent with a role as an electron sink that may help alleviate PSI acceptor-side limitation and rebalance the redox state of the electron transport chain. These findings show that ATP synthase deficiency reorganizes photosynthetic electron flow and promotes prolonged hydrogen production, and point to hydrogenase as a candidate electron sink under energy-constrained conditions.

## Introduction

A delicate fine-tuning of the photosynthetic machinery is required to regulate the electron flux in the thylakoid membrane. A stoichiometric ratio of 3ATP:2NADPH is needed for carbon fixation via the Calvin–Benson–Bassham (CBB) cycle (Rochaix, 2011). Per 2 NADPH formed during linear electron flow (LEF), an insufficient number of protons are pumped into the lumen to produce 3 ATP and fulfill the needs for carbon fixation (Kramer and Evans, 2011). Thus, the extra requirement for ATP is fulfilled by the various processes such as cyclic electron flow (CEF) (Alric, 2010). In CEF a proton gradient is generated by the redirected electron flow and drives ATP synthesis but does not modify the pool of reduced NADPH under illumination, compared to LEF (Joliot and Joliot, 2002). During CEF, electrons are released by PSI and reinjected into the photosynthetic electron transport chain by reducing the plastoquinone (PQ) pool on the stromal side of the thylakoid membrane. In the green microalga *Chlamydomonas reinhardtii* (*C. reinhardtii*) CEF operates through two distinct pathways: a type II NAD(P)H dehydrogenase (NDA2) dependent pathway and a pathway involving proton gradient regulation 5 (PGR5) and PGR5-Like1 (Johnson et al., 2014; Saroussi et al., 2016). In the NDA2 dependent pathway, electrons derived from NAD(P)H are transferred back to plastoquinone (PQ) via the membrane-peripheral NDA2 whereas in the PGR5 pathway, electrons are transferred from ferredoxin back to the PQ pool via a poorly understood ferredoxin–plastoquinone reductase (FQR) (Nawrocki et al., 2019a). The site and identity of this FQR activity remain debated, with competing evidence supporting either activity at cytochrome b₆f complex (Cytb_6_f) or involvement of PGR5/PGRL1 (Buchert et al., 2020; Takahashi, 2022).

The proton motive force (*pmf*) generated by LEF and CEF must be efficiently dissipated to sustain photosynthetic activity. This is achieved by the ATP synthase (cF₁cF_o_) which matches free energies stored in the *pmf* and the phosphorylation potential through its chemical (cF_1_ portion) and electrochemical motors (cF_o_ portion) (Junge and Nelson, 2015). The soluble cF_1_, where ATP is formed from ADP and inorganic phosphate, is mechanically coupled to the membrane-embedded cF_o_, where proton translocation through the a-subunit drives the rotation of the c-ring (Hahn et al., 2018). When cF₁cF_o_ activity is impaired under environmental stress (Kanazawa and Kramer, 2002; Kohzuma et al., 2009; Takagi et al., 2017), proton flux through the enzyme is restricted and the resulting lumen acidification (ΔpH) downregulates the photosynthetic machinery in both PSII and Cytb_6_f. These measures prevent excessive PSI overreduction, which can, in turn, limit light-driven ATP synthesis (Foyer et al., 2012). Specifically, the ΔpH adjusts the fraction of PSII engaged in photochemistry through energy-dependent non-photochemical quenching (NPQ), leading to a downregulation of electron transport and an upregulation of PSII photoprotection (Li et al., 2000). Moreover, the Cytb_6_f is governed by the “photosynthetic control” process where its PQH₂ oxidase activity is slowed under acidic lumen conditions (Rumberg and Siggel, 1969). These conditions also favor state transition NPQ which modulates light harvesting capacity at PSII and PSI. State transitions involve the reversible phosphorylation of mobile antenna complexes that are triggered by the PQ pool redox state (Rochaix, 2014).

In this study, we impose ATP limitations and ΔpH imbalance by cF₁cF_o_ gene lesions (Figure 1), producing photosynthesis-impaired mutants that display extreme photosensitivity under dim light, eventually leading to degradation of the PSII cores and inhibition of electron transfer (Majeran et al., 2001). Moreover, in photosynthetic microbes such as *C. reinhardtii* severe ATP limitation or anaerobic conditions create an energy imbalance that is expected to profoundly influence PSI acceptor-side limitation and hydrogen production under anoxic conditions (Clowez et al., 2015).

**Figure 1:**
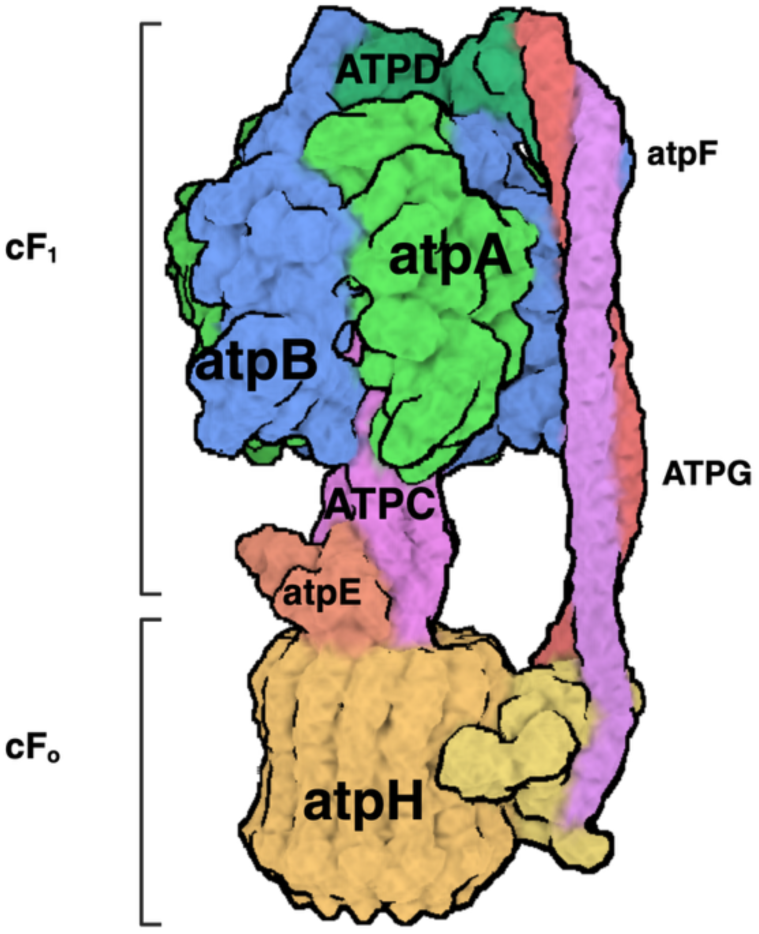
Structure of the chloroplast ATP synthase (cF₁cFo). Subunits are labeled according to their respective gene nomenclatures with nuclear genes in capital letters. Deletions of chloroplast *atpB* (β-subunit) and *atpH* (c-subunit), as well as *ATPC* (γ-subunit) were studied. Created with BioRender.com; PDB accession 6VM1.

In *C. reinhardtii* hydrogenases are expressed under anaerobiosis and work as a safety valve, preventing the photosynthetic machinery from over-reduction (Nagy et al., 2021). Protons are reduced to H_2_ through the chloroplast hydrogenases (HYDA1 and HYDA2) under anoxic conditions (Hemschemeier et al., 2008). Electron transport through PSI via ferredoxin to hydrogenase creates molecular H_2_ that supports the electron transport chain and alleviates PSI acceptor-side limitation under anoxia, when the activation of the CBB cycle is delayed due to limited availability of ATP (Clowez et al., 2015; Godaux et al., 2015). The photoproduction of H_2_ can occur in two processes, direct and indirect. The direct process occurs when photooxidation of water takes place and the PSII and PSI supply reductants to ferredoxin through the electron transfer chain and eventually donated to HYDA1/2 (Melis et al., 2000).

The indirect pathway sources electrons from starch degradation that are buffered in the PQ pool via NDA2 activity (Hemschemeier and Happe, 2005; Posewitz et al., 2009). H_2_ production is limited by the supply of electrons to the hydrogenases because most of the electrons are redirected to ferredoxin-NADP+-reductase (FNR) for carbon assimilation by the CBB (Milrad et al., 2018). Another major challenge for prolonged H_2_ production is HYDA1/2 sensitivity to oxygen (Stripp et al., 2009).

The regulatory responses to ATP limitation within the chloroplast, particularly during the transition to anoxia, remain poorly understood. Earlier studies of ATP synthase mutants focused on their light sensitivity, PSII stability and cyclic electron flow (Lemaire et al., 1988; Majeran et al., 2001; Johnson et al., 2014). Clowez et al. (2015) used the ATP synthase mutant fud50 and hydrogenase-deficient strains to show that, in anoxia, the light-induced reoxidation of PSI acceptors is ATP-dependent and that hydrogenase is the main ATP-independent route adjusting the stromal redox poise. To our knowledge, however, hydrogen production by ATP synthase mutants during illumination in anoxia, and its relation to the electron partitioning of mutants affected in different subunits, has not been characterized. We hypothesized that, when carbon fixation is restricted by the ATP shortage, hydrogenase could act as an alternative electron sink that relieves PSI acceptor-side limitation. To address this, we compared a newly generated chloroplast ΔatpH deletion, the nuclear *atpC1-2* mutant and, for comparison, fud50 by optical spectroscopy and membrane inlet mass spectrometry. We report that the ΔatpH mutant shows strong PSI acceptor-side limitation in oxic cells, which is partially relieved under anoxia and coincides with prolonged hydrogen production.

## Results

The background strain t222 and two cF₁cF_o_ mutants were used in the main figures: the ΔatpH mutant lacks the chloroplast-encoded cF_o_ c-subunit, while *atpC1-2* carries a deletion in the nuclear-encoded cF_1_ γ-subunit.

### ATP synthase deficiency restricts dark relaxation of the electrochromic shift signals

To verify the absence of active cF₁cF_o_ in the mutants we followed electrochromic shift (ECS) decay kinetics. Signals were recorded following single turnover laser flash under oxic and anoxic conditions. The fast phase (“a-phase”; detected at ∼500 µs after the flash) represents the immediate generation of the electric field following charge separation at the photosystems. Next, a slower phase occurred within the millisecond range, corresponding to electron transfer through the Cytb_6_f. The variable decaying phase reflects the consumption of the electrochemical gradient through ionic fluxes, predominantly via cF₁cF_o_ activity. The results (Figure 2 A,C) show that the proton efflux in both cF₁cF_o_ mutants is markedly slower than in t222 under oxic conditions (Figure 2A) and differs under strongly reducing conditions (Figure 2C). Figure 2A shows that both Δ*atpH* and *atpC1-2* display compromised ECS decay due to their mutations. Analysis of the *t*_1/2_ kinetics (Inset tables A,C) reveals that ΔatpH has a significantly slower decay under oxic conditions (725.6 ms) compared to *atpC1-2* (132.26 ms) and t222 (13.85 ms). Following 2 hours of anoxia (Figure 2C), the decay kinetics of *atpC1-2* and t222 slowed even further (154.91ms and 101.2 ms, respectively), whereas the Δ*atpH* had a slightly faster decay (505.44 ms), which remained markedly slower than t222, consistent with persistently limited proton conductance through the thylakoid membrane. The decay kinetics of mutants had a significant difference between genotypes, with no significant main effect of condition (Oxic vs Anoxic), although t222 slowed ∼7-fold under anoxia. Both *atpC1-2* and Δ*atpH* mutants exhibited significantly slower *t*_1/2_ compared to t222 across all conditions.

**Figure 2.**
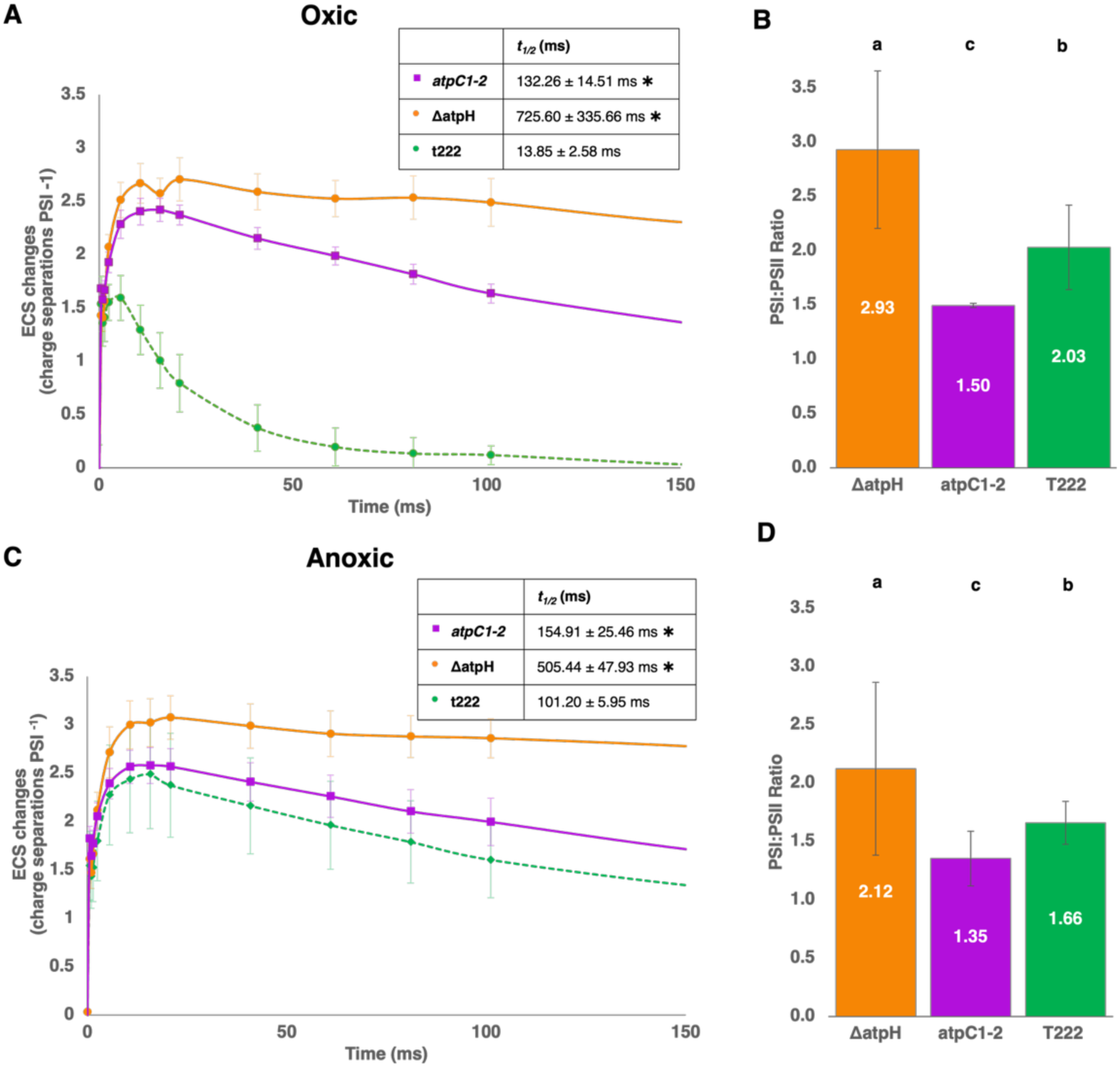
Flash-induced electrochromic shift (ECS) kinetics of ATPase mutants demonstrate an impaired decay of the proton motive force and changes in photosystem ratios. **(A, C)** Dark-adapted cells of ATPase mutants (solid lines) and wild-type (t222: dashed lines), under oxic or anoxic conditions respectively. A laser flash was applied at *t=0* ms and the following ECS kinetics in darkness were monitored. Proton efflux half-times (*t*_1/2_) were determined for each strain under anoxic and oxic conditions. Asterisks (*) indicate statistical significance (*p* < 0.05) relative to the t222 within the same condition, based on Student’s t-tests. Mean values are presented (n ≥ 3). Statistical significance across genotypes was also determined by two-way ANOVA (p<0.0001) followed by post-hoc pairwise comparisons. **(B, D)** The amplitude of the rapid, sub-ms ECS rise following a flash was used to quantify PSI and PSII contributions to the signals in the presence of PSII inhibitors (3-(3,4-dichlorophenyl)-1,1-dimethylurea (DCMU) and hydroxylamine (HA)). The corresponding PSI/PSII activity ratios were calculated, and averaged values are shown (n > 3 ± SEM). Statistical significance between genotypes is denoted by letters (a, b, c) where ’a’ denotes the highest ratio (*ΔatpH*), ’b’ the intermediate (*t222*), and ’c’ the lowest (*atpC1-2*) as determined by Tukey HSD post-hoc test (*p* < 0.05). The statistical grouping is based on a two-way ANOVA.

The ECS assay can also estimate relative contributions of the photosystems. The amplitude of the fast “a-phase” is proportional to the number of active photosystems. By using PSII inhibitors (DCMU and HA), the contribution of each photosystem can be separated: the decrease in signal amplitude after PSII inhibitor’s addition represents the PSII component, while the remaining insensitive fraction corresponds to PSI. Accordingly, the PSII signal was calculated by subtracting the PSI-only signal from the combined (in the absence of inhibitors) PSI+PSII signal, and the PSI:PSII ratio as the PSI-only signal divided by the PSII signal. Under oxic conditions in dark-grown cultures, the t222 wild type showed PSI:PSII ratios of 2.03. This ratio was reduced to 1.66 under anoxia. In Figure 2B Δ*atpH* displayed a higher PSI:PSII ratio (2.93), which could implicate a strong bias toward PSI-driven electron flow associated with a shift in photosynthetic metabolism or an inactivation of PSII. The ratio changed under anoxia (Figure 2D), and Δ*atpH* dropped to 2.12. The *atpC1-2* mutant, on the other hand, maintained low PSI:PSII ratios in both oxic and anoxic conditions (1.50 and 1.35 respectively), suggesting that this mutation imposes a constraint on the photosynthetic system that expresses differently on the PSI:PSII ratios. Genotype had a significant effect on the PSI:PSII ratio, with no significant effect of condition or genotype-condition interaction. For comparison, measurements were also performed on the *fud50* mutant (ΔatpB, lacking the chloroplast-encoded cF₁ β-subunit). This strain exhibited ECS dark relaxation kinetics similar to those observed in the Δ*atpH* and *atpC1-2* mutants, confirming that the restricted proton efflux is a general feature of cF₁cF_o_ deficiency (Supplemental Figure S1).

### Impaired PSII stability and enhanced respiration in the ΔatpH mutant

To evaluate PSII integrity under continuous anoxia, PSII quantum yield (Fv/Fm) was measured using a DUAL-PAM 100 (Figure 3 A,B). Cells were incubated with glucose oxidase, glucose, and catalase to maintain anoxic conditions, Ficoll to prevent sedimentation of the cells and bicarbonate to support PSII stability. Cells were illuminated at 300 µmol photons m⁻² s⁻¹ for 12 h; every hour illumination was interrupted for 10 min of dark relaxation before a saturating pulse was applied to determine Fv/Fm. Residual activity was calculated by normalizing each value to the starting point Fv/Fm measurement (Figure 3A). Upon illumination, the Δ*atpH* mutant exhibited a rapid decline in PSII quantum yield, decreasing by ∼55% relative to the t222 reference (Figure 3A). This difference was statistically significant within the first hour of light treatment. The wild type t222 showed a moderate decline (∼10%) after the first hour, followed by a steady, gradual decrease throughout the remainder of the experiment. The *atpC1-2* mutant followed a similar trend to t222 throughout the experiment with only a non-significant difference in the first 3 h. The *atpC1-2* mutant maintained relatively higher overall yields according to chlorophyll fluorescence measurements; a trend consistent with ECS-based PSII estimations from Figure 2. The relatively low initial Fv/Fm values may be attributed to dark-induced State 2 transition in TAP medium (Bulté et al., 1990; Nawrocki et al., 2019b). The lack of recovery (Figure 3B) after dark relaxation may point to sustained PSII impairment in the Δ*atpH* mutant, with Fv/Fm dropping from 0.305 to 0.055. While t222 shows a continuous mild decline over the 12-h time course (Figure 3A), it recovered after 1 h of dark relaxation to Fv/Fm levels comparable to those at the beginning of the experiment, whereas ΔatpH did not (Figure 3B). *atpC1-2* preserved most of its photosynthetic yield throughout the experiment. The difference in recovery across strains was significant, highlighting that *atpC1-2* retains substantial photosynthetic stability despite defected ATP synthase function, whereas ΔatpH is particularly sensitive to prolonged illumination.

**Figure 3.**
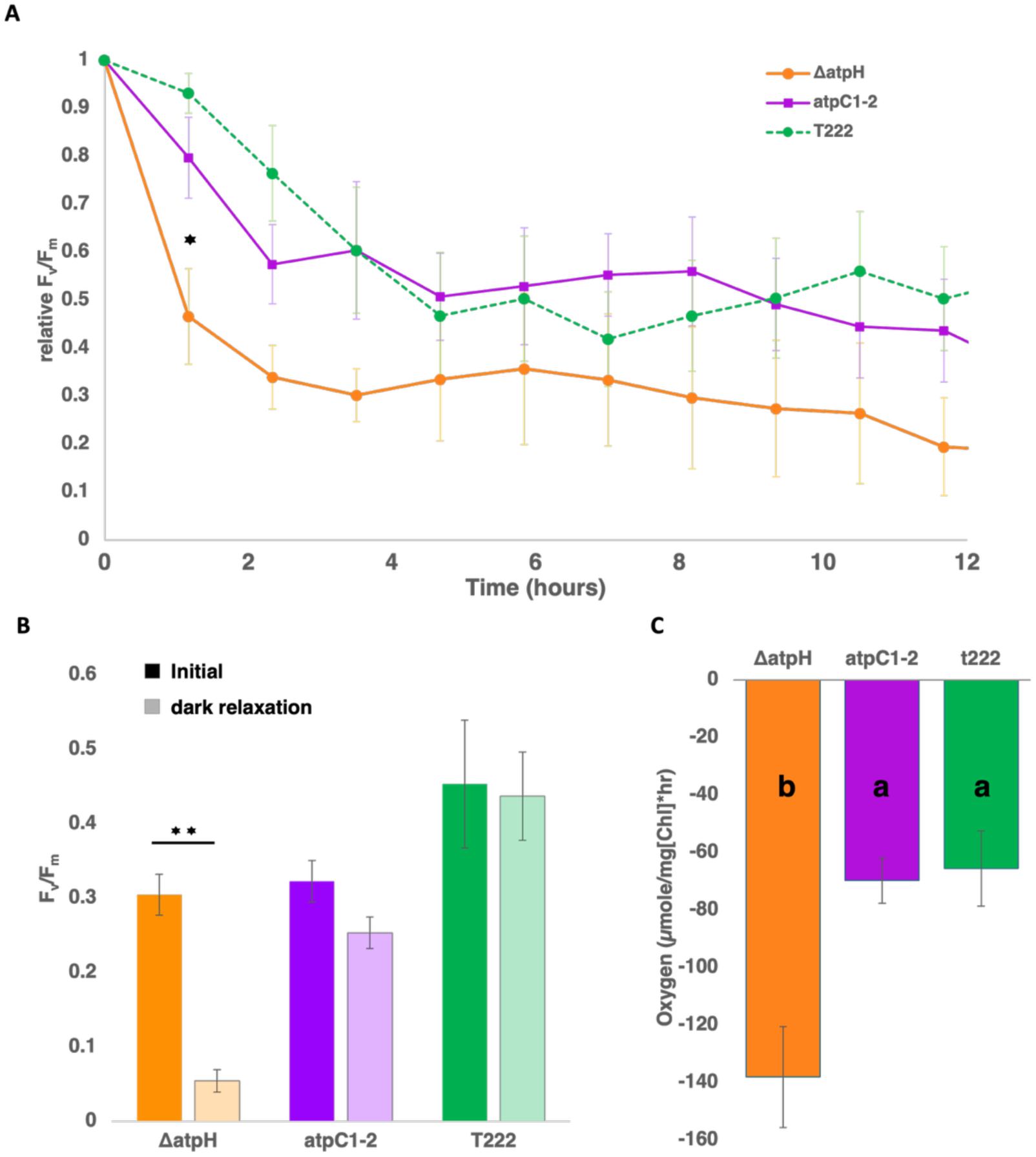
ATPase deficiency lowers PSII efficiency and increases respiration in ΔatpH. **(A)** Change in maximum PSII quantum yield monitored over 12 hours of illumination (interrupted hourly by 10 min of dark relaxation for each measurement; 300 μmol photons m^-^² s^-^¹). Data are normalized to the initial dark-adapted state. Data are presented as the mean ± SEM from n>4 biological replicates. Asterisks indicate significant differences from wild type (Welch’s *t*-test *p=0.026). **(B)** Initial Fv /Fm (solid bars) was measured prior to light exposure, and post-illumination recovery (light bars) was determined after 1 h dark relaxation. Values represent mean ± SEM from n>4 biological replicates. Statistical differences between the initial and recovery states were determined by paired two-tailed *t*-tests (**p = 0.0027). A two-way ANOVA confirmed a significant strain × timepoint interaction (p=0.0134). **(C)** Dark oxygen uptake rates in the mutant strains compared to the parental line. Letters within the bars indicate statistical groupings, different letters represent significant differences at p<0.05 based on a one-way ANOVA followed by Tukey’s HSD post-hoc test. The overall difference among the groups was statistically significant (One-way ANOVA, p = 0.015; Tukey’s HSD, p = 0.021).

To assess metabolic performance of the respiratory chain, dark oxygen consumption rates were measured to determine the respiratory capacity of each strain (Figure 3C). The Δ*atpH* mutant showed a significant 2-fold change in respiration compared to its background strain t222 (137.9±17.56 and 65.48±13.13 µmol O₂ (mg Chl)⁻¹ h⁻¹, respectively). The *atpC1-2* mutant showed no significant difference from the wild type (69.7 ± 7.78). These results show elevated respiration in the Δ*atpH* mutant, suggesting an increased mitochondrial activity to support the chloroplast ATP pool.

When evaluating the results, one must consider the genetic background of the *atpC1-2* strain. The atpC1-2 strain originates from the cell-wall-deficient atpC1 mutant (Drapier et al., 2007) and was backcrossed into a cell-wall-containing background. The *atpC1-2::ATPC* rescue strain was therefore included in the experiments to confirm whether the phenotypes resulted from ATP synthase deficiency. Taken together, cF₁cF_o_ impairment elicits metabolic rewiring in a genetic background-dependent manner, tested at the levels of dark respiration and PSII photochemistry under light exposure.

To evaluate growth capacity, biomass accumulation was monitored under mixotrophic conditions (Supplemental Figure S2). The *atpC1-2* mutant maintained a growth profile comparable to the WT (t222) while the rescue strain (rescue *atpC*1-2::*ATPC*) reached the highest final density. The Δ*atpH* strain exhibited a more pronounced impairment in biomass accumulation. The Δ*atpH* mutant grew at a slower rate throughout the exponential phase, failing to reach wild-type density. At the end of the log phase, Δ*atpH* displayed a decrease in optical density. This suggests that even low-light settings eventually caused inhibitory effects in this cF₁cF_o_ mutant strain.

We next focused on PSI and determined electron transfer rates to monitor consequences of ATP shortage in the plastids.

### Altered PSI redox states and cyclic electron flow

To further understand the electron partitioning, we measured P700^+^ redox states before and after the addition of PSII inhibitors, to distinguish PSI performance under full capacity and CEF only conditions in absence of PSII. (Figure 4 A,B) presents a simplified quantification of P700 redox components YI (PSI effective yield), YNA (acceptor-side limitation), and YND (donor-side limitation) highlighting distinct redox behaviors among the cF₁cF_o_ mutants.

**Figure 4.**
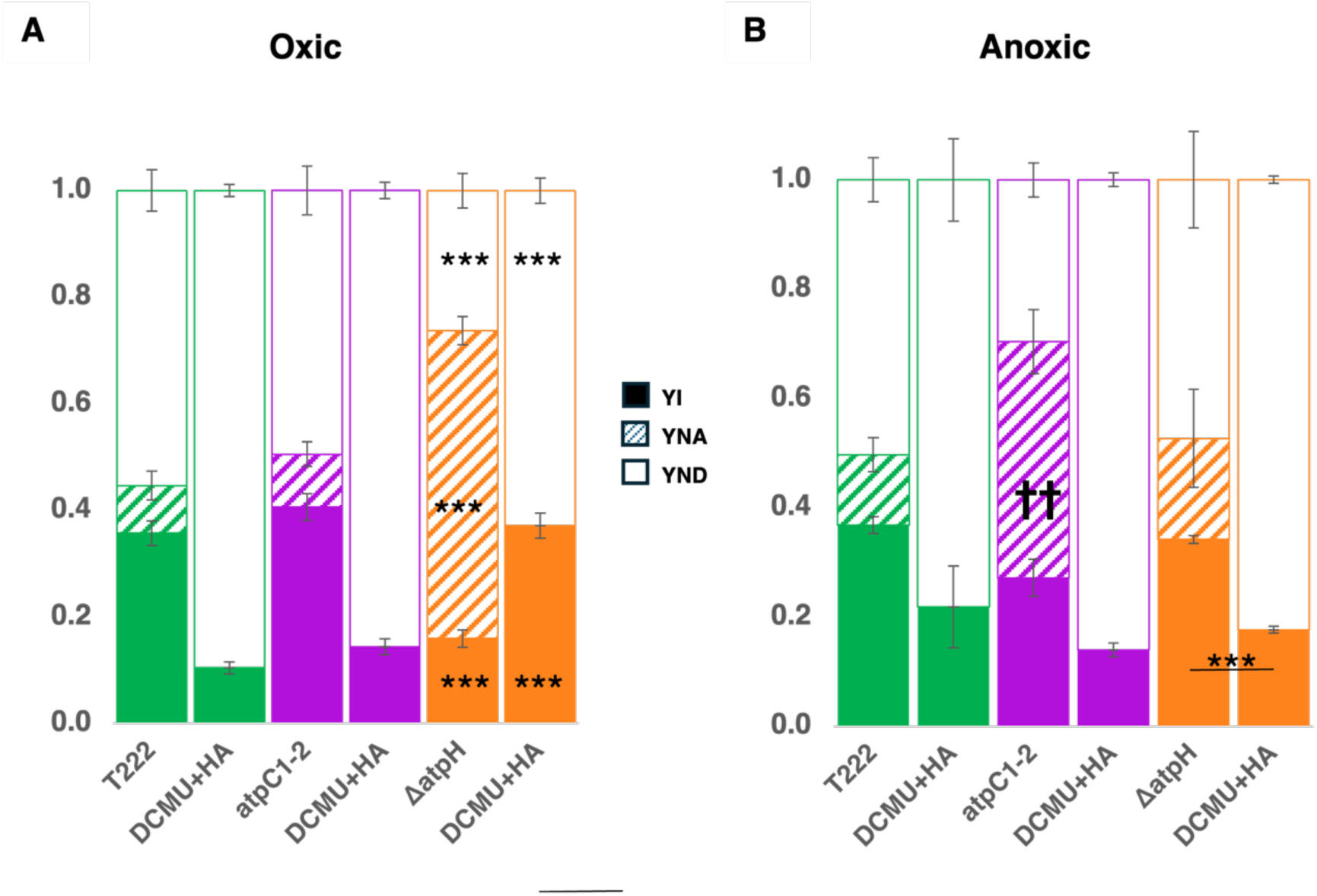
P700 redox states of cF₁cFo mutants and WT under oxic and anoxic conditions. P700 dynamics were recorded in cF₁cFo mutants (*atpC1-2*: purple, Δ*atpH*: orange), and WT (green: t222), PSI parameters measured under oxic **(A)** and anoxic **(B)** conditions are shown (mean ± SEM, n ≥ 3 biological replicates). PSI parameters comprise quantum yield (YI, filled bars), acceptor-side limitation (YNA, striped bars), and donor-side limitation (YND, open bars). Asterisks inside bar segments indicate statistically significant differences relative to the wild-type (t222) under identical conditions (∗∗∗p<0.001, Welch’s t-test). Daggers inside bar segments indicate a significant change between oxic to anoxic state of the same strain (††p<0.01). The horizontal line with asterisks in (B) indicates significance upon inhibitor addition within each strain (∗∗∗p<0.001).

In oxic conditions, the *atpC1-2* strain exhibited YI and YND values comparable to the WT t222, with only minor YNA contributions. In contrast, the Δ*atpH* mutant displayed a distinctly elevated YNA and a corresponding smaller YND (*p* < 0.001 vs. WT for both), indicating a severe acceptor side limitation under oxic conditions. Upon the addition of PSII inhibitors, which abolishes LEF, YNA was largely absent in all strains. In the Δ*atpH* mutant, this treatment resulted in a recovery of YI to levels comparable to those measured for the parental strain without inhibitors, accompanied by an increase in YND; this donor side limitation was significantly lower than that observed in WT with inhibitors (*p* < 0.001). The relatively high YI under inhibited conditions (*p* < 0.001 vs. treated t222) may indicate a higher CEF capacity where a significant fraction of P700 remains redox active, which will be quantified at the end of this section. Combined with the ECS ratios (Figure 2B), these kinetics suggest that the PSI core in ΔatpH remains functional but is subject to a severe acceptor side bottleneck during linear electron flow, likely stemming from limited ATP availability.

Under anoxic conditions in Figure 4B, the *atpC1-2* mutant displayed an increased YNA (compared to its oxic state *p* = 0.005), indicating an enhanced acceptor side limitation accompanied by a decrease in both YI and YND, while the WT remained largely unchanged across conditions. Upon the addition of PSII inhibitors, the overall trends resembled those observed under oxic conditions, where YNA was largely abolished and YND increased correspondingly. However, compared to oxic DCMU conditions, a slight increase in YI was observed for both t222 and *atpC1-2* under anoxia. The ΔatpH mutant strayed from this pattern, showing a significantly smaller YNA (*p* = 0.028) together with an increase in both YI (*p* < 0.001) and YND upon transition to anoxic conditions. Following the addition of PSII inhibitors, ΔatpH displayed a significantly lower YI (0.177 vs. 0.342, *p* < 0.001), with YND constituting the dominant component of the redox partitioning.

P700^+^ reduction kinetics were measured concurrently with ECS under PSII inhibition to evaluate the turnover efficiency of the cyclic pathway (Table 1). CEF was isolated, and the rates were normalized to ECS-based PSI activity following the framework of (Takahashi et al., 2013). The ECS rise kinetics used for Table 1 are provided in Supplemental Figure S3. The ΔatpH mutant displayed higher rates under oxic conditions (209.26), but rates similar to WT under anoxia (104.86), suggesting a high capacity for PSI-driven cyclic flow. The *atpC1-2* mutant showed only slight differences between conditions (53.09 and 52.28, respectively). In the WT t222, CEF increased significantly under anoxic conditions, a ∼2.4-fold shift from 42 to 100.86 PSI e^-^s^-1^, as expected based on previous reports (Alric, 2010; Johnson and Alric, 2013; Joliot et al., 2022). By contrast, the ΔatpH mutant appears to be shifted excessively toward CEF even under oxic conditions. Upon transition to anoxia, a significant downregulation of CEF was observed in the ΔatpH mutant, whereas *atpC1-2* exhibited no significant change between conditions. In the WT CC-125, CEF likewise increased under anoxia (from 104.85 to 186.77 e⁻ s⁻¹ per PSI), whereas the fud50 (ΔatpB) mutant showed rates similar to CC-125 under oxic conditions (114.15) and lower rates under anoxia (144.53; Supplemental Figure S3F). Under anoxia, alternative electron sinks become available, and electrons may instead be redirected to hydrogenase rather than being used in a futile attempt to pump additional protons into the lumen in the absence of ATP synthase. Because these CEF measurements were performed with PSII inhibited, a condition under which hydrogen production is strongly reduced (Supplemental Figure S4), this possibility is addressed only indirectly in the next section, where hydrogen production is monitored directly.

**Table 1.**
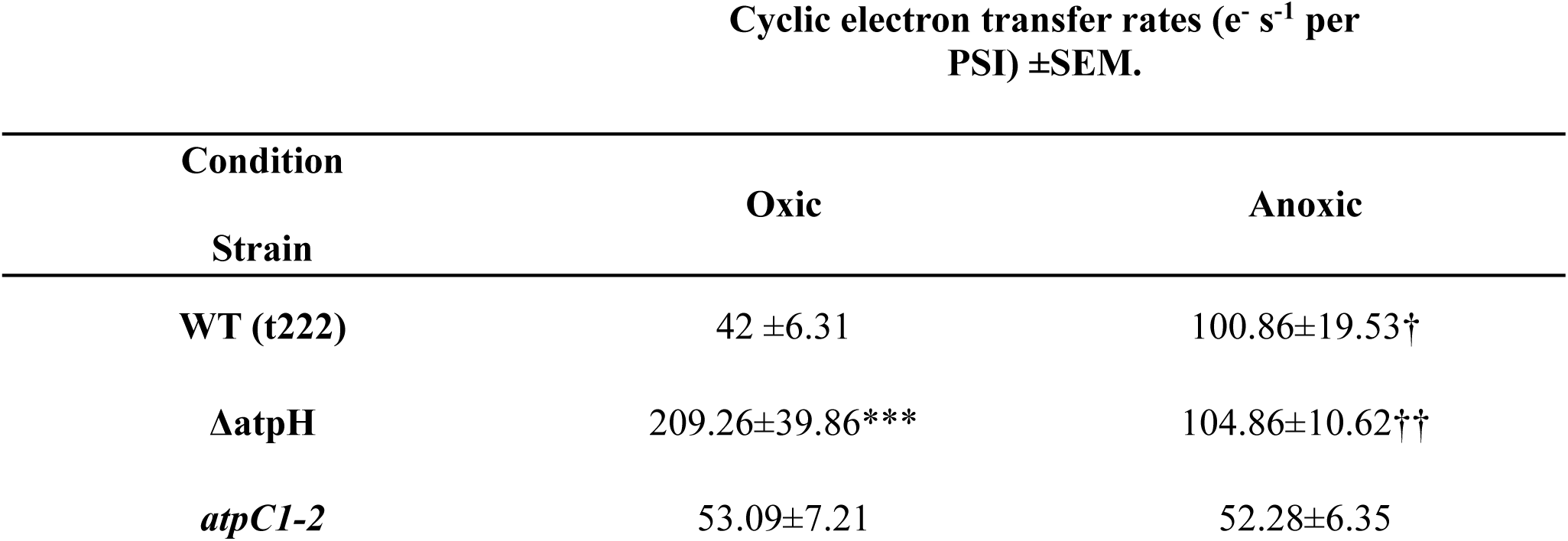
Cyclic electron transfer rates in wild type and mutant strains in oxic and anoxic conditions. CEF rates were determined following the method described in (Takahashi et al., 2013) in the presence of DCMU and HA to inhibit LEF. Data represent the average of at least 3 independent experiments ± SEM. Statistical significance was determined using a Two-Way ANOVA followed by a Tukey post-hoc test. Asterisks indicate significant differences relative to WT (t222) under the same condition (\*\*\**p* < 0.001). Daggers signify a significant change upon transition from oxic to anoxic in the same strain ( †*p* < 0.05, ††*p* < 0.01).

### Hydrogen and CO₂ production in cF₁cF_o_ mutants (MIMS analysis)

Hydrogen and CO₂ exchange were monitored by membrane inlet mass spectrometry (MIMS) after 2 h of dark incubation to establish anaerobic conditions (Figure 5). Illumination at 357 µmol photons m^-^² s^-^¹ triggered photosynthetic H₂ production in all strains. Experiments were performed in TAP medium, as the mutants cannot grow photo-autotrophically and acetate was supplied to provide reducing equivalents for mitochondrial respiration.

**Figure 5.**
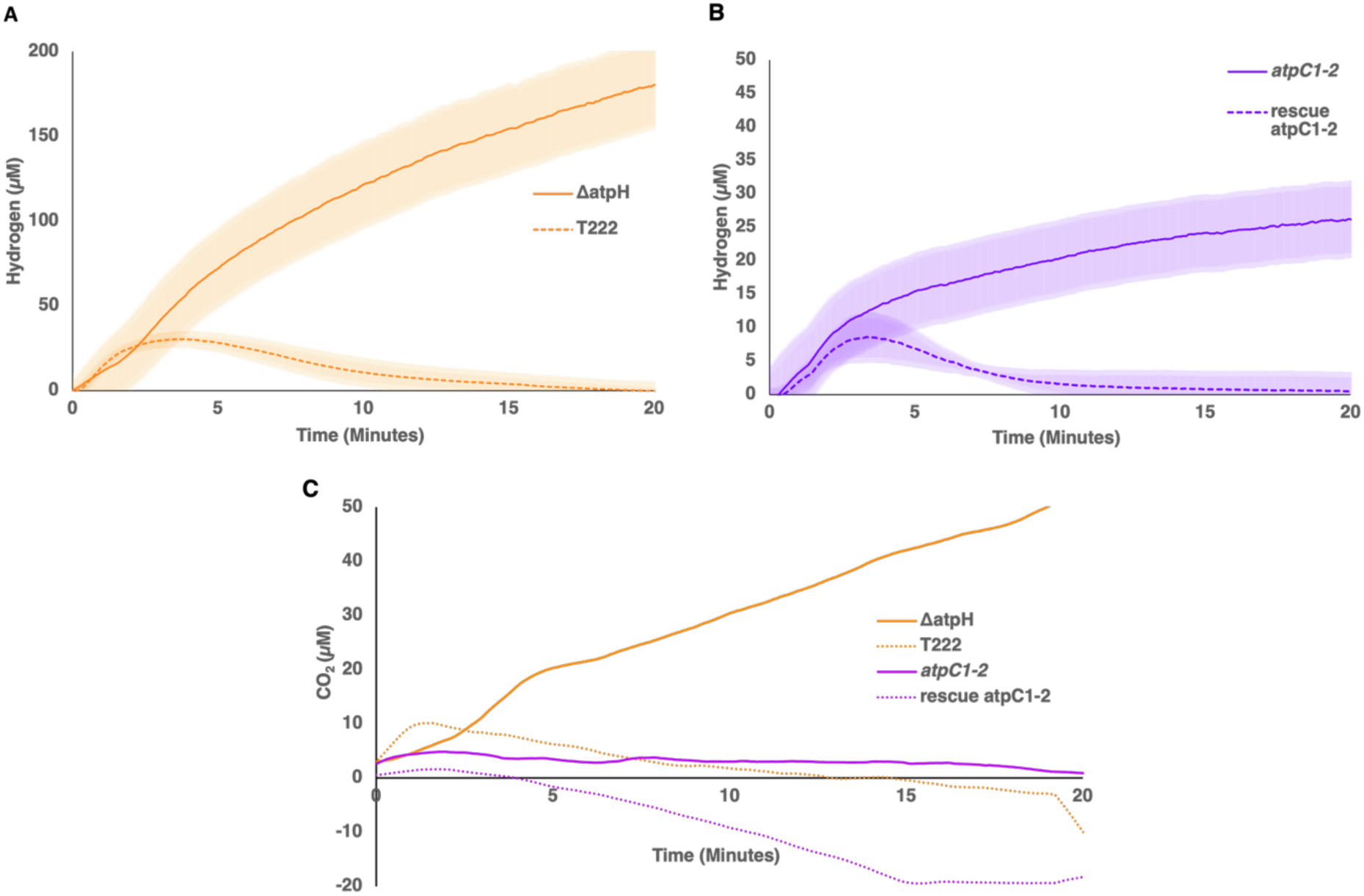
The ATP synthase mutants present a prolonged H2 production compared to their rescues and WT. H2 and CO2 gas accumulation were measured by MIMS. **(A)** ΔatpH and t222 as wild type control. **(B)** *atpC1-2* and its rescue. Cells were brought to a concentration of 20 µg Chl mL^-1^ and were incubated in the dark for 2 h to induce anaerobiosis, after which they were exposed to 20 min of constant illumination (357 µmol photons m^-2^ s^-1^). Shown are the averaged H2 accumulation concentrations. Shaded areas represent the standard error of the mean (SEM), calculated from n>3 independent biological replicates. **(C)** CO2 exchange was monitored for all strains and performed simultaneously to the H2 measurements under the same conditions. Data represent the mean of n>4 biological replicates.

The Δ*atpH* mutant sustained the highest H₂ production, reaching ∼200 µM, comparable to previously reported levels in the pgr5 mutant (Elman et al., 2022), surpassing the WT t222, which ceased H₂ production within 2 minutes of illumination (Figure 5A). The *atpC1-2* mutant also maintained H₂ production, albeit at lower levels than Δ*atpH* (∼30 µM). In the *atpC1-2* rescue strain, H₂ production ceased after ∼3 minutes (Figure 5B). Accompanying the hydrogen-producing phenotype, the Δ*atpH* mutant exhibited a steady increase in CO₂ levels whereas the *atpC1-2* showed a steady trace consistent with respiratory CO₂ release balancing residual fixation. The rescue strain and WT showed a decrease indicative of carbon fixation (Figure 5C). The gradual CO₂ rise in ΔatpH and the lack of net CO₂ uptake in *atpC1-2* are consistent with respiratory CO₂ release exceeding CO₂ fixation, which net gas exchange cannot separate. This outcome is expected if the mutants fail to generate sufficient ATP to sustain the ATP:NADPH ratio required for Calvin cycle activation, which would leave the cycle restricted and could divert electrons to HYDA1/2 for H₂ production under anoxic conditions. To determine whether electrons supporting hydrogen production originate from PSII via the direct pathway or from indirect metabolic routes, H₂ evolution rates were measured by MIMS in the presence and absence of PSII inhibitors (DCMU and HA). Inhibition of PSII strongly minimized H₂ production in all strains, indicating that LEF is the primary source of electrons for hydrogen evolution (Supplemental Figure S4). In the t222 wild-type strain, PSII inhibition completely abolished H_2_ production, whereas the *atpC1-2* mutant retained a small residual activity. In contrast, the Δ*atpH* mutant maintained a detectable fraction of H_2_ production even under PSII inhibition, suggesting that a portion of the electrons is supplied via an indirect pathway.

The *fud50* strain, lacking the cF₁ β-subunit (ΔatpB; Lemaire et al., 1988), also produced H₂ continuously for ∼20 minutes, reaching approximately 90 µM, accompanied by gradual CO₂ accumulation, indicating limited carbon fixation. The corresponding WT exhibited only transient H₂ evolution, which ceased after ∼3 minutes of illumination (Supplemental Figure S5). In the atpB rescue strain, H₂ evolution was again only transient and CO₂ levels decreased, as in the wild type (Supplemental Figure S5), which links the fud50 phenotype to the loss of atpB. Taken together, these traits provide a favorable environment for sustained hydrogen production: limited carbon fixation, which in wild-type cells outcompetes hydrogenase for electrons (Milrad et al., 2018), and, in ΔatpH, elevated respiration that may help to keep oxygen low during illumination. Consequently, electrons may be redirected toward HYDA1/2, which would then serve as a major electron sink under these conditions.

## Discussion

In this study, we examined various cF₁cF_o_ mutations expected to severely limit ATP synthesis in the chloroplast, and, possibly as a coping mechanism for this energetic stress, we observed a partial relief of PSI acceptor-side limitation under anoxia in ΔatpH. This response was accompanied by prolonged H₂ production, which was also seen in *atpC1-2* and fud50. It is important to consider that the strains in this study were generated using distinct genetic approaches and their distinct age span is more than four decades: Δ*atpH* was produced via chloroplast direct knockout via homologous recombination of t222 for this study, while *atpC1-2* was generated through nuclear transformation of a cell wall-deficient cw-15 strain (Smart and Selman, 1991) that was further backcrossed into a cell wall-containing background (Drapier et al., 2007). The *fud50* was generated as a mutation arising from homologous recombination in the chloroplast genome (Woessner et al., 1984). Thus, a degree of background variation between these lines must be acknowledged, which may contribute to the observed baseline differences in their physiological responses. The ECS-based PSI:PSII ratios also differ between the laboratory WT strains t222 and CC-125 (cf. Figure 2 and Supplemental Figure S1), indicating background-specific phenotypes.

Our flash ECS results suggest that the ‘laboratory age’ of the three cF₁cF_o_ mutants may contribute to our experimental readouts, given some striking similarities of the most ancient *fud50* with Δ*atpH* (cf. Figure 2 and Supplemental Figure S1), although isogenic strains would be required to test this. Yet, there are some parameters that differ among the cF₁cF_o_ mutants: When compared with Δ*atpH*, the ECS decay slowdown was substantially less pronounced in *atpC1-2* (Figure 2), which could reflect a different expression of cF₁cF_o_-independent ion fluxes. The latter might also account for P700 redox state differences among both cF₁cF_o_ mutants, owing to distinct ΔpH control of LEF and CEF in oxic vs. anoxic conditions (Figure 4). The impact on PSII was also less dramatic in *atpC1-2* (Figure 3) which, together with its partial survival in the absence of fixed carbon and WT-like mixotrophic growth (Supplemental Figure S2), points to an attenuated damage to the *atpC1-2* chloroplast through mutant-specific recruitments of chemical energy from cellular bioenergetic pathways. Importantly, it is known that second-site mutation(s) can restore photosynthesis in cF₁cF_o_ mutants (Lemaire et al., 1988), and further tests may reveal the specific genetic basis behind the unexpected *atpC1-2* growth phenotype.

Initially, PSII was suspected to be particularly vulnerable, given the mutants’ increased light sensitivity during mixotrophic growth (Supplemental Figure S2) and the anticipated photoinhibitory effects of an elevated ΔpH, which can destabilize the PSII oxygen evolving complex (Krieger and Weis, 1993; Majeran et al., 2001; Johnson et al., 2014). Moreover, it has been proposed that ATP availability is essential for PSII repair, as insufficient ATP prevents the FtsH protease from mediating D1 protein turnover, thereby predisposing PSII to photoinhibition (Preiss et al., 2001; Nixon et al., 2010; Malnoë et al., 2014). Yet, our results demonstrate that PSII remains functionally intact under dark growth with an impairment apparent only after the onset of illumination (Figure 3). Consistent with this, PSII-dependent water oxidation remains the predominant electron source for H_2_ production in the cF₁cF_o_ mutants (Supplemental Figure S4). A smaller fraction of the reductant may originate from pyruvate oxidation by pyruvate:ferredoxin oxidoreductase, which generates reduced ferredoxin during anoxic fermentation and may become increasingly relevant under severe ATP limitation (Chochois et al., 2009; Catalanotti et al., 2013; Milrad et al., 2021).

Despite this initially preserved activity, PSII performance declines during prolonged illumination. After one hour, Fv/Fm, measured following 10 min of dark adaptation, decreases rapidly in the Δ*atpH* mutant (Figure 3A). This decrease may reflect photoinhibition caused by the adverse effect of excessive ΔpH on PSII stability (Krieger and Weis, 1993). Alternatively, it could partly result from enhanced state transition NPQ associated with an over-reduced PQ.

As chloroplast ATP synthesis is compromised, increased reliance on mitochondrial metabolism may help maintain cellular energy balance through ATP exchange between the two organelles (Lemaire et al., 1988; Cardol et al., 2009). Consistent with this interpretation, the Δ*atpH* mutant displayed elevated respiration rates, indicative of enhanced chloroplast-mitochondrial metabolic coupling under stress (Johnson et al., 2014; Bailleul et al., 2015; Elman et al., 2022; Peltier et al., 2024). High respiration rates observed in the Δ*atpH* mutant (Figures 3C, 5C) may contribute to conditions favorable for hydrogenase activity by counterbalancing PSII-derived O_2_ evolution and maintaining hypoxia in the cell. This is particularly important given the oxygen sensitivity of the hydrogenase catalytic site (Stripp et al., 2009). However, respiration alone is unlikely to account for sustained H_2_ production, as the *atpC1-2* mutant also exhibits prolonged H_2_ evolution despite respiration rates comparable to WT, which has an inherently high respiration rate (Figure 3C).

The impaired cF₁cF_o_ function appears to impose broad constraints on photosynthetic efficiency, producing a phenotype at the PSII core that is reminiscent of sulfur deprivation. In *Chlamydomonas*, extended H_2_ production is commonly induced by sulfur deprivation, which inhibits D1 turnover and thereby causes the progressive loss of PSII activity and oxygen evolution (Tsygankov et al., 2002; Nagy et al., 2018). The prolonged H_2_ production observed in the cF₁cF_o_ mutants (Figure 5A, B) might therefore be related to a comparable compromise in PSII function, triggered intrinsically by defective ATP synthesis rather than by nutrient deprivation, although our Fv/Fm measurements, which span hours, cannot show that PSII loss underlies the H₂ phenotype in the first 20 min.

This bioenergetic reorganization also strongly affects carbon metabolism. The mutants show no net CO_2_ assimilation, while Δ*atpH* even exhibits net CO_2_ accumulation (Figure 5C), consistent with a severe restriction of CBB cycle activity (net CO₂ exchange also includes respiratory release), most likely owing to insufficient plastidic ATP availability. This raises the question of how the mutants meet their remaining chloroplast ATP demand. One possible route is the import of phosphorylated carbon intermediates through triose-phosphate or related phosphate translocators, which could support an indirect ATP shuttle between acetate-fuelled mitochondria and the chloroplast (Heldt and Rapley, 1970; Johnson and Alric, 2013; Huang et al., 2023). Increased CEF is another typical response to an unfavorable ATP:NADPH balance, as it redirects electron transport toward additional *pmf* generation without producing NADPH (Joliot and Joliot, 2002; Rott et al., 2011). Although CEF is enhanced in ΔatpH under oxic conditions (Table 1) but not in the other cF₁cF_o_ mutants, it cannot fully compensate for defective photophosphorylation because the resulting *pmf* cannot be efficiently converted into ATP.

Recent analyses of a “broken Calvin-Benson cycle” mutant of *C. reinhardtii*, ΔrbcL, revealed a shift from phototrophic toward predominantly heterotrophic metabolism (Saint-Sorny et al., 2022). Given the divergent responses of PSI (Figure 4) and PSII (Figure 3) in these “broken ATP synthesis” mutants, similarly comprehensive analyses could help resolve their distinct strategies for utilizing light energy and the accompanying metabolic rewiring across cellular compartments. It is conceivable that under anoxic conditions, non-cyclic processes such as electron diversion to hydrogenase may compete with the FQR pathway.

Our data are correlative, and two interpretations cannot yet be distinguished. In the first, hydrogenase acts as an electron sink that relieves PSI acceptor-side limitation. In the second, the limited carbon fixation (which in wild-type cells outcompetes hydrogenase for electrons; Milrad et al., 2018) leaves hydrogenase as the main remaining sink for the residual electron flow from PSII and indirect sources, without hydrogenase being required to relieve PSI limitation. The CEF measurements were performed in the presence of DCMU and HA, conditions under which hydrogen production is strongly diminished (Supplemental Figure S4), so the lower CEF of ΔatpH under anoxia cannot be attributed to electron diversion to hydrogenase on the basis of the present data. Distinguishing between these possibilities will require hydrogenase-deficient backgrounds (for example, hydA1 hydA2 mutants combined with the chloroplast ΔatpH lesion by crossing), in vivo P700 measurements during hydrogen evolution, and direct measurements of ATP, pmf and oxygen exchange. Finally, the three mutants differ in genetic background and age, and only ΔatpH and fud50, both chloroplast-encoded deletions, show comparable ECS relaxation and H₂ production, whereas the elevated oxic CEF of ΔatpH was not observed in fud50 (Supplemental Figure S3F); conclusions on the general consequences of ATP synthase deficiency should therefore be drawn with caution.

In these mutants, the energetic stress imposed by ATP limitation may be partially alleviated by hydrogenase, which could act as a safety valve that diverts excess electrons and thereby relieves redox pressure on the photosynthetic apparatus. Hydrogen production provides a qualitative readout of electron flow under anoxic conditions and can therefore serve as a proxy for electron partitioning within the chloroplast. The coincidence of H₂ production with a partial relief of PSI acceptor-side limitation in ΔatpH under anoxia is consistent with our data. Whereas the mutant is severely constrained under oxic conditions, anoxia may provide an alternative exit for electrons through hydrogen evolution. This suggests that for Δ*atpH*, anoxia initially represents a state of relative relief rather than increased stress, albeit prolonged illumination eventually leads to PSII photoinhibition (Figure 3A). The phenotypic variance observed between the *atpC1-2* strain and the Δ*atpH* mutant highlights a range of metabolic flexibility.

The differing degrees to which these strains partition electrons into the hydrogenase sink suggest that structural disruption to the cF₁cF_o_ complex can promote H_2_ production to some extent. This difference may have a biotechnological implication: partial impairment of the cF₁cF_o_ complex could be explored as a way to modulate H₂ production. However, the mutants pay a considerable cost in growth and PSII stability (Figure 3; Supplemental Figure S2), which would have to be overcome in any engineering strategy.

## Materials and methods

### Plasmid construction and chloroplast transformation

The *cemA*/*atpH* region (starting with the *cemA* CDS and extending 732 bp downstream of *atpH*) was amplified from WT chloroplast DNA using restriction site-containing primers 1 (*Eco24I*) and 2 (*BamHI*) (for oligonucleotide sequences, see Supplemental Table S1). The *Eco24I*/*BamHI*-digested PCR product was ligated into a similarly digested pUC18 cloning vector and propagated as pUC18-*cemA*_*atpH* in NEB 5-alpha *E. coli* cells using ampicillin selection. In the next step, the aminoglycoside adenyltransferase (aadA) spectinomycin-resistance cassette (Fu et al., 2017) was amplified from a pUC18-aadA vector using phosphorylated primers 3 and 4. Double digestion of pUC18-*cemA*_*atpH* with *ClaI* and *EcoRV* removed a 307-bp fragment containing *atpH*. The 5′ overhang generated by *ClaI* was blunted using T4 DNA polymerase, and the vector was subsequently dephosphorylated with FastAP. The phosphorylated aadA PCR product generated with primers 3 and 4 was then ligated into the linearized vector, resulting in recircularization and replacement of *atpH* with the selectable marker cassette, yielding plasmid pUC18-Δ*atpH*. Mutant strains were generated by chloroplast transformation and selection on spectinomycin (150 µg mL⁻¹). The WT strain t222 was used as recipient. Spectinomycin-resistant transformants were repeatedly replated to obtain homoplastic lines. Homoplasmy was confirmed by PCR using primers 5 and 6, which yielded the expected 2.35-kb Δ*atpH*::aadA fragment and no detectable 1.23-kb WT fragment.

### Growth conditions

*Chlamydomonas reinhardtii* strains (wild-types t222 and CC-125, mutants Δ*atpH*, *atpC1-2*, fud50 (ΔatpB), an atpB rescue line, and the rescue line *atpC1-2*::*ATPC* (Buchert et al., 2017)) were grown heterotrophically in TAP medium at 24°C. The cells were grown in the dark. Cultures were kept in Erlenmeyer flasks under constant stirring and continuous darkness. Prior to physiological measurements, cells were diluted with fresh TAP medium to reach the early-log phase (2–5 µg Chl mL⁻¹). Total chlorophyll was extracted in 80% (v/v) acetone and quantified according to (Jeffrey and Humphrey, 1975).

Cells were cultivated in a Multi-Cultivator OD-1000 system (Photon Systems Instruments, Brno, Czech Republic) in 80 mL tubes with continuous aeration. Growth curves of wild-type and mutant strains were assessed in TAP medium under continuous illumination (30 μmol photons m⁻²s⁻¹) at 24 °C. Cultures were inoculated at an initial optical density of OD₇₂₀ 0.1 and maintained for 5 days, with OD_720_ recorded at 10-minute intervals to monitor biomass accumulation. For spot assays, cells were spotted at an initial OD_750_ of 0.1 onto TAP (mixotrophic) or TP (photoautotrophic) agar plates and incubated under continuous illumination (30 µmol photons m⁻² s⁻¹) or in the dark.

### MIMS Gas-Exchange Measurements (H_2_, O_2_ and CO_2_)

To monitor real-time gas exchange, cells were centrifuged for 5 min at 4000 rpm at room temperature and resuspended in TAP medium (50 mM HEPES) to a final chlorophyll concentration of 20 µg Chl mL⁻¹. The suspension was placed in a 5 mL sealed Starna quartz cuvette (Starna Cells) and subjected to a 2-hour dark incubation under constant stirring to reach anoxia. Following incubation, the cuvette was fitted into a metabolic chamber (optical unit ED-101US/MD, Walz) and exposed to actinic light at an intensity of 357 µmol photons m⁻² s⁻¹ using a Dual-Pulse Amplitude Modulated Fluorometer (DUAL-PAM-100, Heinz Walz GmbH, Effeltrich, Germany). Where indicated, DCMU (10 µM) and HA (1 mM) were added 10 minutes prior to light exposure to inhibit specific pathways. Mass signals for H_2_, CO_2_, ^16^O_2_ were recorded with a 0.5-s dwelling time per mass. Raw traces were converted to gas exchange rates using conversion factors derived from a standard curve, as previously described by (Liran et al., 2016). A detailed explanation how to perform calculation appears in Supplemental Methods.

### Oxygen measurements

Dark respiration rates and oxygen consumption were recorded using an optical sensor (FireSting, PyroScience) equipped with an OXROB3 probe. Cells were harvested by centrifugation for 5 min at 4000 rpm at room temperature and resuspended in TAP medium (50 mM HEPES) to a final concentration of 10 µg Chl mL⁻¹. The cell suspension was transferred to a 4 mL respiration vial (OXVIAL4, PyroScience) and maintained under complete darkness with constant stirring. Oxygen uptake was calculated via linear regression analysis of the oxygen curves and normalized to the total chlorophyll concentration of the sample.

### DUAL-PAM 100

Chlorophyll fluorescence was recorded using a DUAL-PAM-100 fluorometer (Heinz Walz GmbH, Effeltrich, Germany). For anoxic measurements, cells were supplemented with glucose oxidase (200 U mL⁻¹), catalase (200 U mL⁻¹) and glucose (50 mM) to remove oxygen, 10% (w/v) Ficoll to prevent sedimentation, and 2 mM bicarbonate to support PSII stability. Cells were harvested by centrifugation and resuspended in fresh media to a final density of 10 μg Chl mL⁻¹. Actinic red light (300 μmol photons m⁻² s⁻¹) was provided by DUAL-DR modules, with intensity calibrated using a US-SQS/L light detector and a Li-250A light meter (LI-COR Biosciences). To assess photosynthetic stability, the maximum quantum yield of PSII (Fv/Fm) was monitored over a 12-hour period under constant actinic illumination. Measurements were performed at 1-hour intervals, where each saturating pulse was preceded by a 10-minute dark relaxation period to ensure the reopening of PSII reaction centers and the relaxation of non-photochemical quenching. Fv/Fm was calculated as (Fm − F0)/Fm using the saturation pulse method at each time point.

### Optical spectroscopy

Optical spectroscopy measurements were carried out by using a Joliot-type spectrophotometer (JTS-150, SpectroLogiX), using cells that were adapted to alternating 2.5 s darkness and 2.5 s actinic light (AL; 550 µmol photons m^-2^ s^-1^ peaking at 630 nm). Briefly, electron transport rates (ETR) were obtained in the presence of LEF but referred to PSI charge separation activity. This stems from separate ECS calibration measurements that allowed, via saturating single-turnover flashes, for the conversion of optical changes (obtained as Δ*I*/*I* at 520 nm - 546 nm) into the ECS signals corresponding to 1 PSI charge separation (Bailleul et al., 2010). The ETR protocol was based on a Dark Interval Relaxation Kinetics approach (Sacksteder and Kramer, 2000; Joliot and Joliot, 2002). P700 measurements relied on the Klughammer & Schreiber method (Klughammer and Schreiber, 1994).

### Electrochromic shift

In accordance with (Buchert et al., 2020; Lebok and Buchert, 2024), *in vivo* changes in the electric field component of the *pmf* across the thylakoid membrane were monitored using the electrochromic shift (ECS). Cells were centrifuged at 5000 rpm and resuspended in TAP with 10% Ficoll to prevent cell sedimentation. The sample was adjusted to a final concentration of 20 µg Chl mL⁻¹ and 5 mL was placed within JTS-150 spectrometer operated in dark-pulse absorbance mode. For the PSI:PSII ratios, PSII was inhibited with DCMU (10 µM) and HA (1 mM). For anoxic measurements, cells were incubated in the dark for 2 hours under constant stirring. A detailed JTS script as well as graphical explanations are provided (Supplemental Figure S6) as well as explanation how to perform calculations appear in Supplemental Methods and Supplemental Tables S2, S4 and S5. Pulsed detection light was provided by LEDs peaking at 520 nm and 546 nm, driven at a probe-beam current of 76,734 µA with ultra-short 2 µs detecting pulses to optimize the signal-to-noise ratio without inducing unintended actinic pre-illumination. The light-detecting diodes were protected from scattered actinic light by 3-mm BG39 filters (Schott, Mainz, Germany). Charge separation was induced by a single-turnover laser flash generated by a DCM dye laser pumped by a frequency-doubled Nd:YAG laser (Litron Nano). ECS kinetics were monitored using repeated dark-interval relaxation kinetics (DIRK) measurements under steady-state actinic illumination. The complete acquisition sequence is described in Supplemental Methods and schematically illustrated in Supplemental Figure S6.

### P700

P700 redox kinetics were measured using a JTS-150 LED pump–probe spectrometer (SpectroLogiX). Samples were exposed to continuous actinic illumination and periodically probed using dark-interval measurements. Maximal P700 oxidation was induced by saturating pulses, and subsequent reduction kinetics were followed over several seconds using logarithmically distributed acquisition schedules. Detailed acquisition timing is provided in Supplemental Figure S7 and Supplemental Methods (detection settings and data processing in Supplemental Tables S3–S5). Cells were harvested by centrifugation (5000 rpm) and resuspended in TAP medium supplemented with 10% Ficoll to minimize sedimentation during measurements. Chlorophyll concentration was adjusted to 20 µg chlorophyll mL^-^¹. For anoxia, cells were incubated in the dark for 2 hours under constant stirring. Absorbance was monitored using pulsed detection LEDs centered at 705 and 740 nm in combination with interference filters (FWHM 10 and 15 nm, respectively). The difference signal ΔI/I_705-740_ was used to follow P700 oxidation and reduction. Scattered actinic light was minimized using RG695 cut-off filters (3 mm thickness; Schott, Mainz, Germany) placed in front of the photodiodes. Steady-state P700 oxidation under continuous actinic illumination was defined as P_s_. Maximal P700 oxidation induced by a saturating pulse was defined as P_m_. To determine the absolute maximal oxidizable P700 pool (P^’^_m_), samples were incubated with hydroxylamine (1 mM) and DCMU (10 µM) to completely block electron donation from Photosystem II.

The effective quantum yield of PSI (YI) was calculated as:

(Pm — Ps)/P’m.

Acceptor-side limitation was calculated as:

(P’m — Pm)/P’m.

and donor-side limitation as:

Ps/P’m.

Technical replicates were averaged prior to analysis. Signals were baseline-corrected using measurements acquired immediately after the time-axis reset (average of the ten pre-illumination points; see Supplemental Methods).

### P700^+^ Relaxation Kinetics and CEF Calculation

Cyclic electron flow (CEF) was estimated according to the approach described by Alric (2010) and Takahashi et al. (2013). Under steady-state illumination, the redox state of P700 reflects a dynamic equilibrium between photooxidation of PSI reaction centers and their re-reduction by electrons returning through the photosynthetic electron transport chain. At steady state, the rate of P700 oxidation is equal to the rate of P700 reduction (k_ox_ ⋅[P700_red_]) = (k_red_ ⋅[P700_ox_]). To isolate the cyclic component of electron transport, linear electron flow from Photosystem II was blocked by addition of 10 µM DCMU and 1 mM hydroxylamine (HA). Under these conditions, electrons reaching PSI originate predominantly from cyclic pathways.

The apparent rate constant of P700 photooxidation (k_ox_) was determined independently from electrochromic shift (ECS) measurements (Supplemental Figure S3). This parameter reflects the effective photochemical turnover rate of PSI under the applied illumination conditions and depends on both incident light intensity and the functional antenna size of PSI.

The fraction of reduced P700 ([P700red]) during steady-state illumination was determined using the saturation-pulse method (Klughammer and Schreiber, 1994). Briefly, the additional P700 oxidation induced by a saturating pulse was used to quantify the fraction of PSI centers that remained available for oxidation immediately before the pulse.

The electron flux through PSI was then calculated as: k_ox_ ⋅ [P700_red_] This equation represents the rate of electron transfer through PSI. Under conditions where electron donation from PSII was abolished, PSI electron flux provides a direct estimate of cyclic electron flow around PSI (Alric, 2010).

### Statistics

Spectroscopic and kinetic measurements were performed using 3-7 independent biological replicates (n = 3-7) and are reported as mean ± SEM. For multi-parameter calculations, standard errors were propagated using Gaussian error propagation. Differences between genotypes and oxygen conditions (oxic vs. anoxic), including genotype × condition interactions, were evaluated using two-way ANOVA followed by Tukey’s multiple-comparisons test or Welch’s t-test. Student’s t-tests, paired two-tailed t-tests, and one-way ANOVA followed by Tukey’s HSD test were used where indicated in the figure legends; the number of biological replicates (n) is given in each figure legend. Statistical significance was defined as p < 0.05. Kinetic fitting and statistical analyses were performed in Python using the SciPy library.

## Acknowledgments

We would like to thank Dr. Michael Vaughn for helpful discussion in support with JTS experiments. We would like to Thank the Yacoby lab members for their help during the study.

## Funding

This study was financed by grants from the Israel Science Foundation 941/22, Ministry of Science and Technology (MOST) grant 8395 and the Council for Higher Education (VATAT) Excellence Center - Energy. F.B. acknowledges funding from the Deutsche Forschungsgemeinschaft (project 507704013 as part of DFG FOR 5573/1, and project 461765884).

## Conflicts of Interest

The authors declare no conflict of interest.

## Data Availability Statement

All data supporting the findings of this study are available in the Supplemental Data; the raw data files are provided as additional Supplemental Data files.

## Author contributions

J.R.M. conceived the study, designed and performed experiments, analyzed the data, and wrote the manuscript. T.E. provided guidance and assistance throughout the entire study. F.B assisted with JTS-based measurements, contributed to data interpretation and manuscript editing. I.Y. conceived the study, supervised the project, contributed to experimental design, wrote the manuscript, and secured funding.

## Supplemental Information

**Supplemental Table S1.**
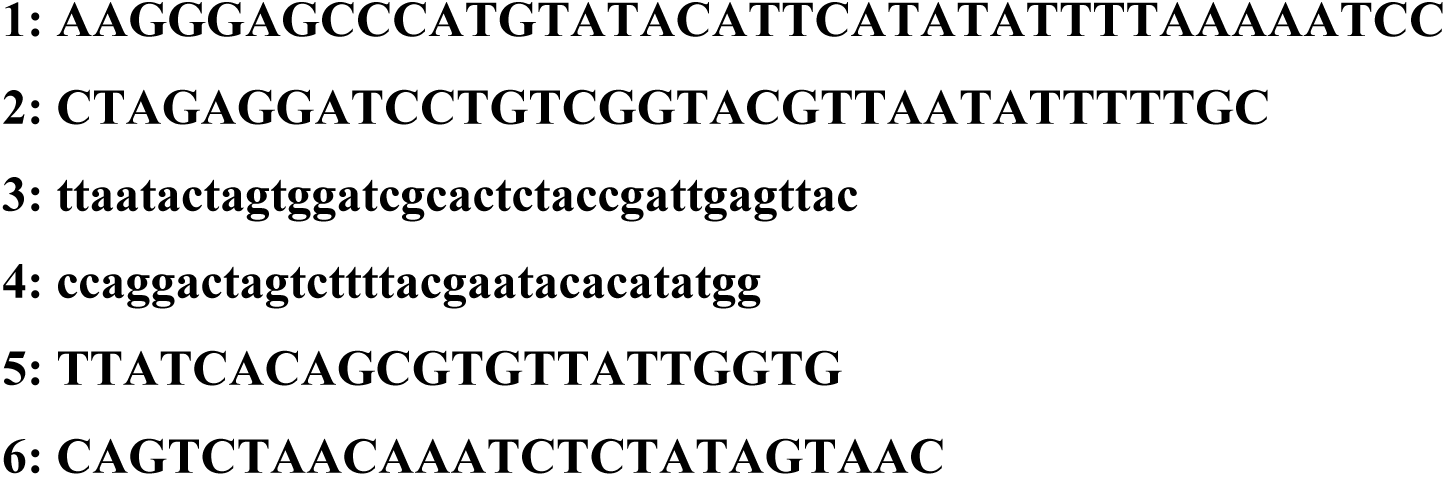
Oligonucleotide sequences.

**Supplemental Figure S1.**
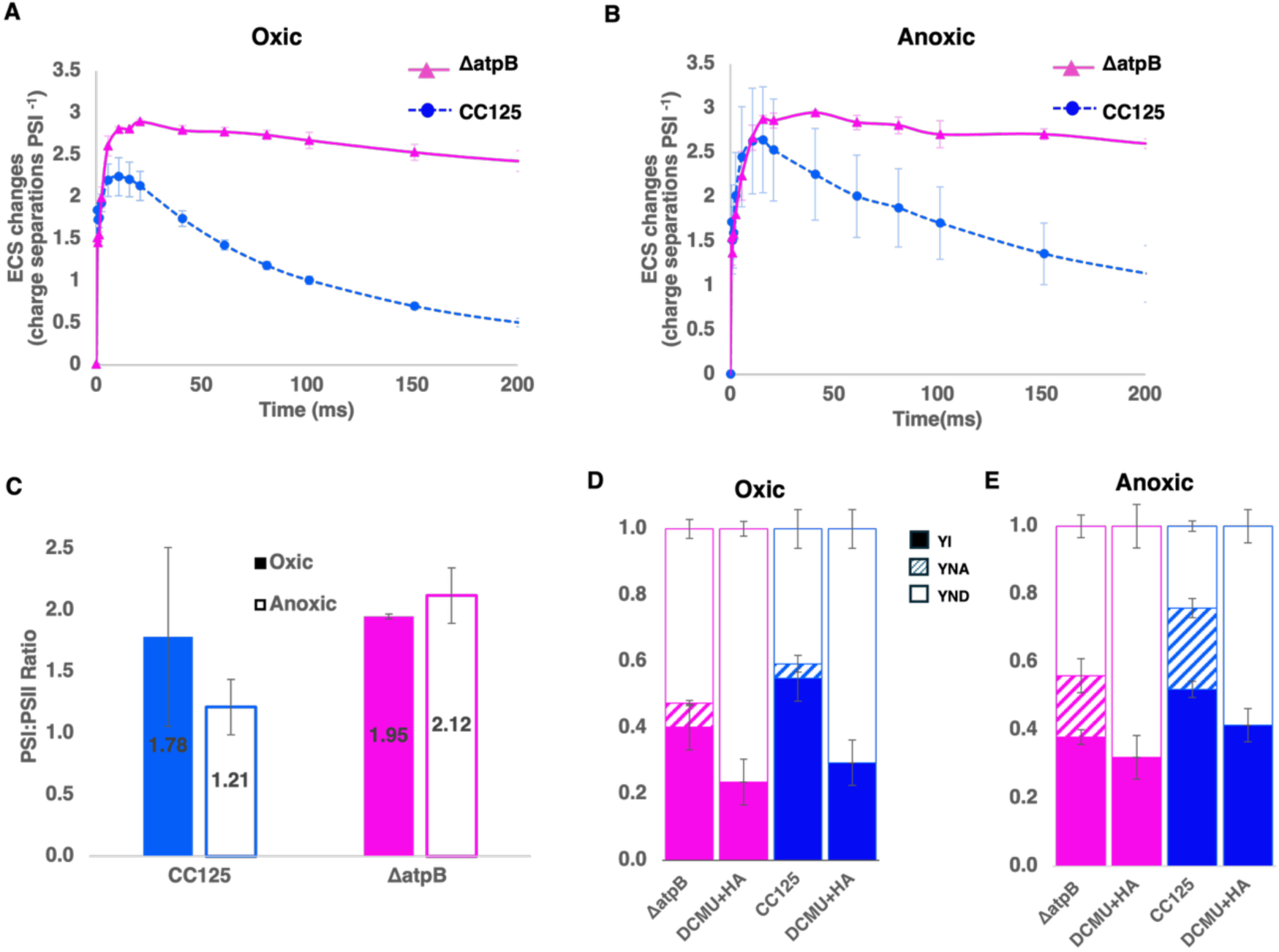
ECS and P700 kinetics of ΔatpB mutant and WT strain CC-125. (**A, B**) ECS kinetics of dark-adapted cells following a short laser flash at *t* = 0 s, monitored as absorbance changes at 520-546 nm, under oxic (**A**) and anoxic (**B**) conditions. (**C**) PSI:PSII activity ratios derived from ECS measurements in the presence of the PSII inhibitors DCMU and HA. (**D,E**) P700 parameters measured under oxic (**D**) and anoxic (**E**) conditions, including PSI quantum yield (YI, filled bars), acceptor side limitation (YNA, striped bars), and donor side limitation (YND, open bars). Data are presented as mean ± SEM (*n* > 3).

**Supplemental Figure S2.**
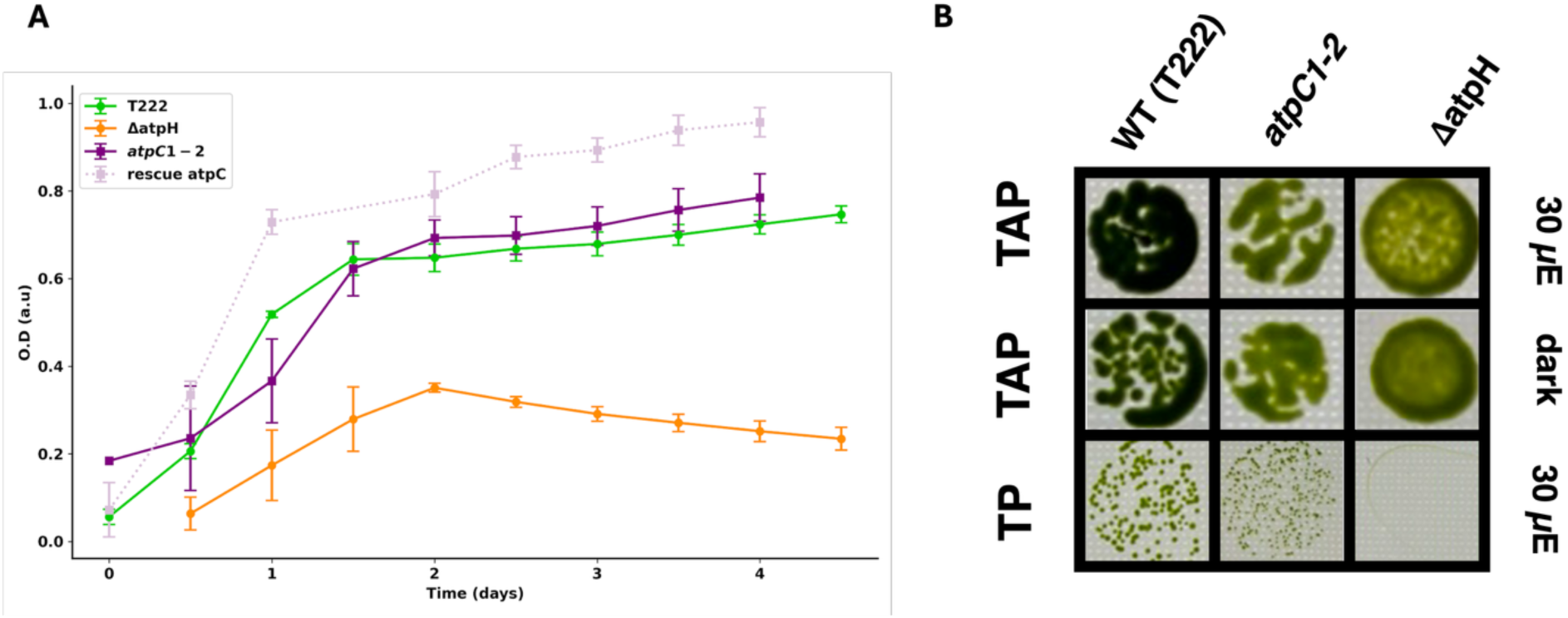
Growth analysis of wild type and ATP synthase mutant strains. **(A)** Growth curves under mixotrophic conditions. Optical density (OD720) was monitored online for 5 days using TAP medium at an illumination of 30 μmol photons m^-2^ s^-1^ (initial OD720 = 0.1). Data points represent the mean of four independent biological replicates ± standard error (SE) for t222 (green), ΔatpH (orange), *atpC1-2* (purple), and the atpC1-2::ATPC rescue strain (lilac, dotted line). **(B)** Spot assays on solid media. *C. reinhardtii* strains (t222, *atpC1-2*, and ΔatpH) were spotted at an initial cell density of OD750 =0.1 onto TAP (mixotrophic) or TP (photoautotrophic) agar plates. Plates were incubated under either continuous illumination or dark conditions to assess phenotypic growth differences. Representative images of three technical replicates are shown.

**Supplemental Figure S3.**
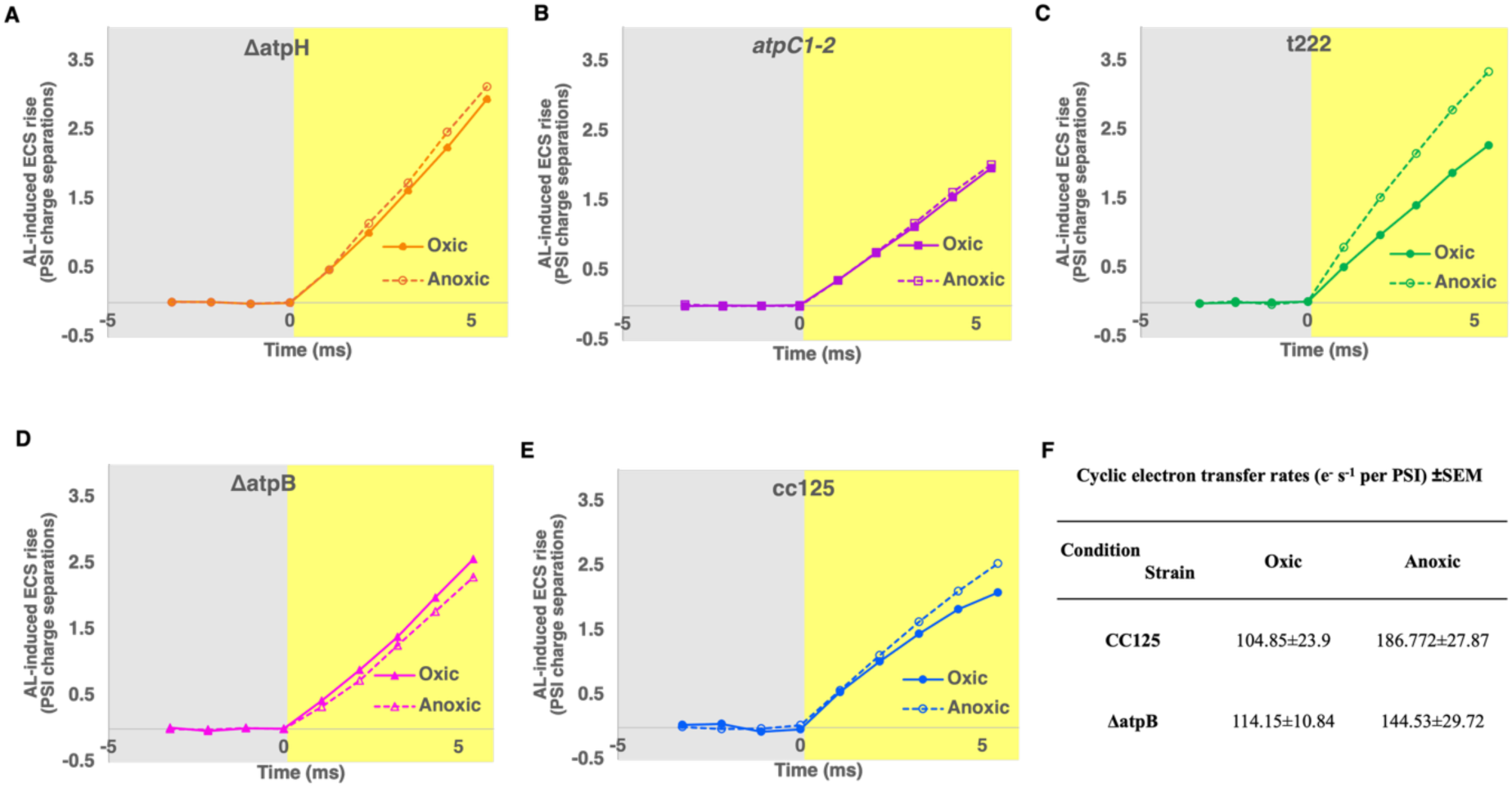
Initial actinic light-induced electrochromic shift (ECS) kinetics in wild-type and ATP synthase mutant strains under oxic and anoxic conditions used for CEF calculations. Initial ECS kinetics are shown at the onset of actinic light in the presence PSII inhibitors to isolate cyclic electron flow. ECS signals were normalized to the ECS amplitude generated by a single-turnover saturating flash, the light-induced ECS rise is expressed as PSI charge separations. The initial linear ECS slope and the PSI photochemical yield Y(I) obtained from P700 measurements were used to calculate the rate of cyclic electron flow. **(A)** ΔatpH, **(B)** *atpC1-2* **(C)** t222 **(D)** ΔatpB and **(E)** CC-125 under oxic and anoxic conditions. Filled symbols and solid lines represent oxic conditions, open symbols and dashed lines represent anoxic conditions. Gray shaded regions indicate the dark period and the yellow - the onset of actinic light. Data are presented as the mean from n>3 independent biological replicates. **(F)** CEF rates derived from the corresponding electron transport measurements, with LEF inhibited by DCMU and HA, under oxic and anoxic conditions.

**Supplemental Figure S4.**
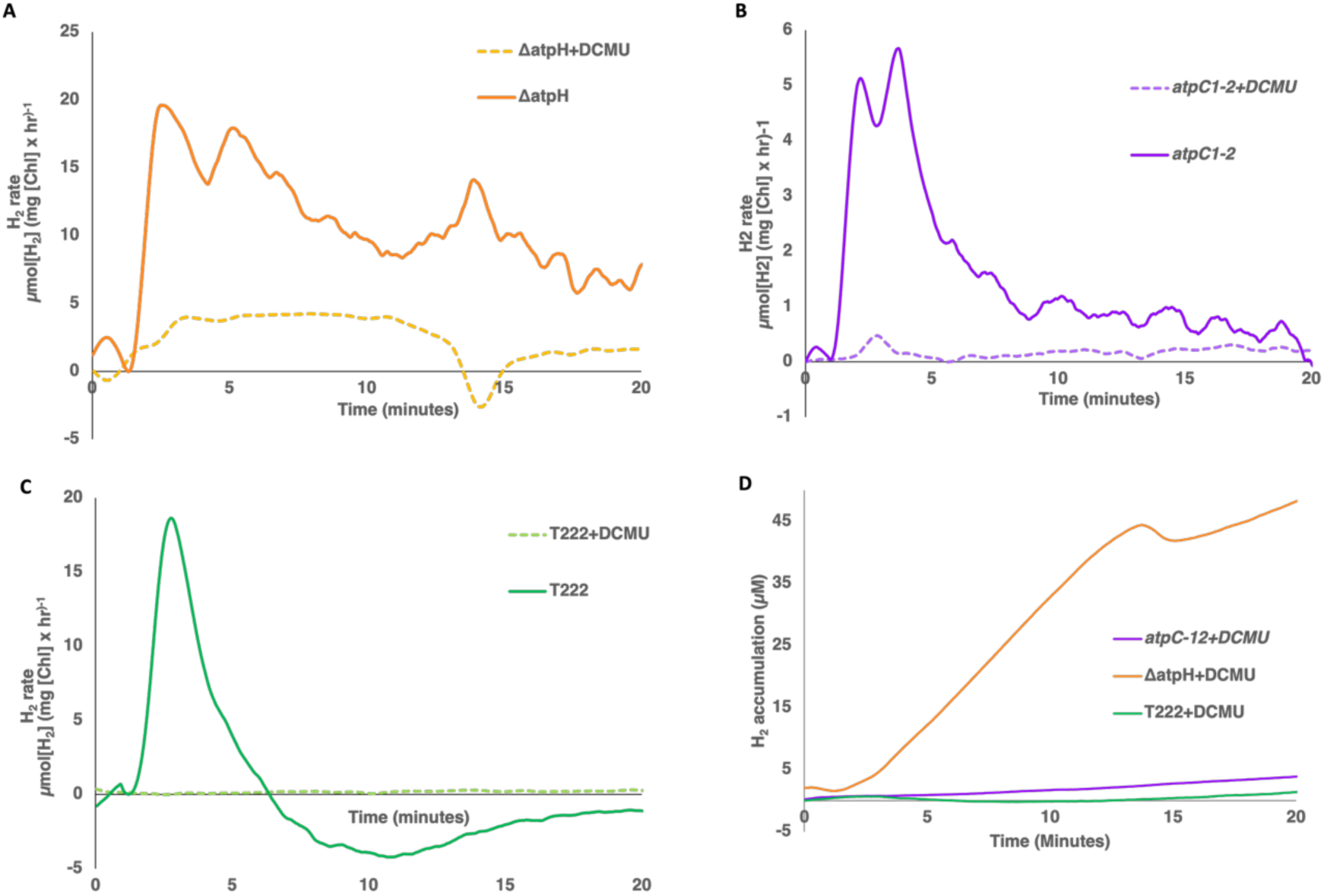
MIMS analysis of direct and indirect hydrogen production reveals functional linear electron flow and PSII activity under anoxic conditions. H₂ production rates were measured by MIMS. Illumination at 357 µmol photons m⁻² s⁻¹ was applied at t = 0, while inhibitors were added 10 min prior to illumination. H2 production rates in the presence and absence of inhibitors for (A) ΔatpH, (B) *atpC1-2*, and (C) t222 wild type (direct rates calculated as total minus indirect pathway). PSII was inhibited by the addition of DCMU (10 µM) and hydroxylamine (HA, 1 mM). (D) Total H2 accumulation comparison among all strains.

**Supplemental Figure S5.**
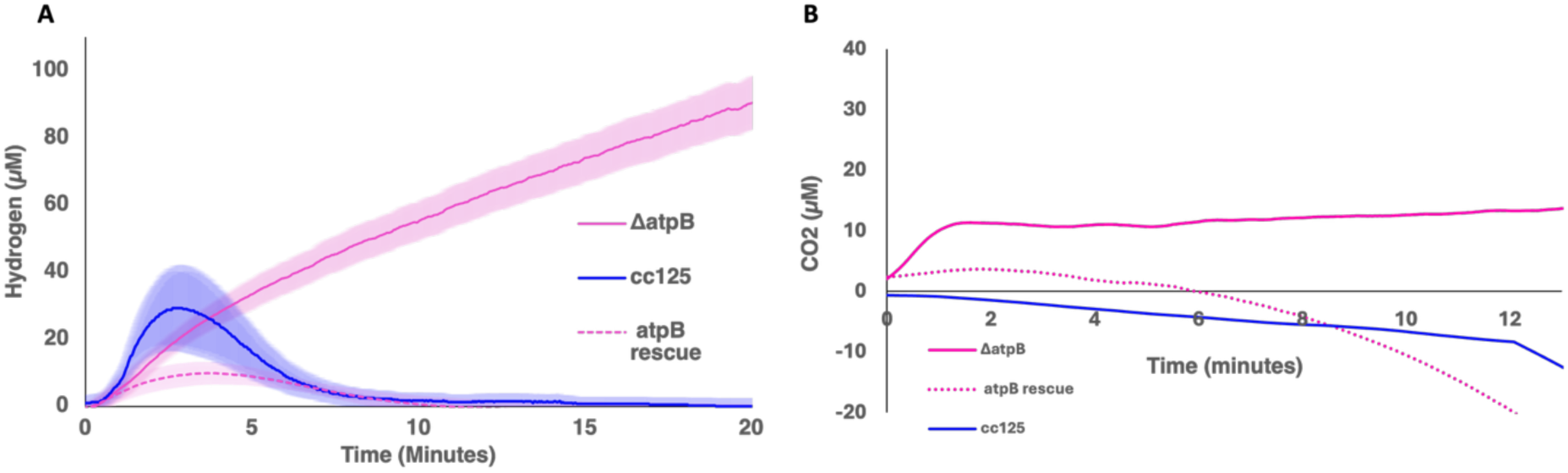
The fud50 (ΔatpB) mutant presents a prolonged hydrogen production compared to WT. H2 and CO2 gas accumulation were measured by MIMS. **(A)** ΔatpB, its WT background CC-125 and the atpB rescue strain were measured under anaerobiosis and exposed to 20 min of constant illumination (357 µmol photons m^-2^ s^-1^). Shown are the averaged H2 concentrations of n>3 biological repeats of each strain. Shaded areas represent the standard error of the mean (SEM). (**B)** CO2 exchange was monitored for all strains and performed simultaneously to the H2 measurements under the same conditions. Data represent the mean of n>4 biological replicates.

## Supplemental Methods

**Supplemental Figure S6.**
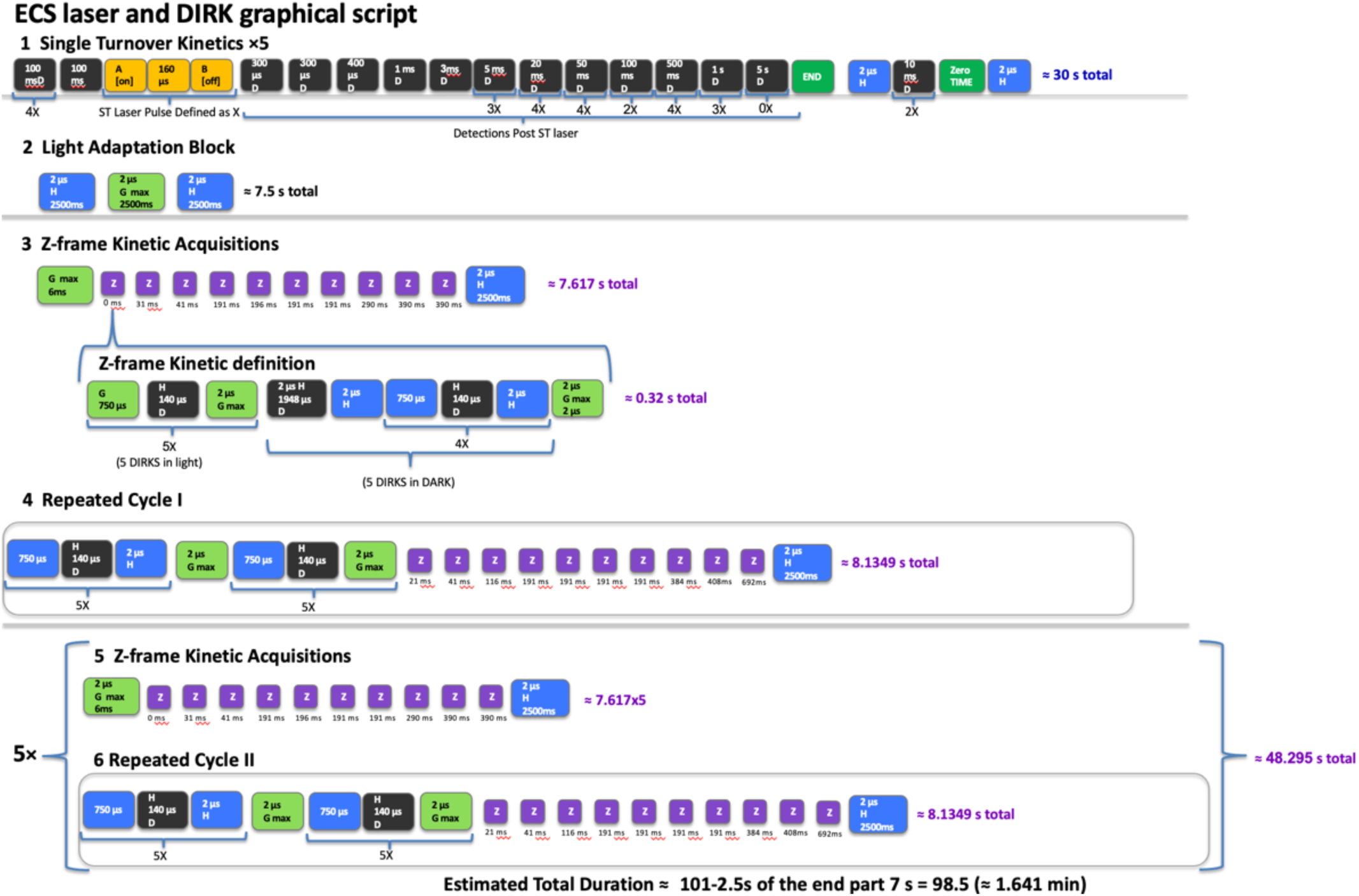
Graphical representation of the JTS script for electrochromic shift (ECS) and dark-interval relaxation kinetics (DIRK) measurements. (1) Single Turnover Kinetics: Tracks the rapid movement of electrons across the membrane induced by a single light flash followed by a dark recovery phase. (2) Light Adaptation Block: Exposes the cells to consecutive phases of darkness, continuous light, and a post-illumination baseline to adapt the sample. (3) Z-frame Kinetic Acquisitions: Measures rapid light and dark micro-cycles to capture membrane potential changes under steady-state conditions. (4) Repeated Cycle I: Alternates brief light and dark windows to monitor the initial phase of photosynthetic induction. (5, 6) Repeated Cycle II: Repeats the measurement sequence across five consecutive cycles to capture long-term performance and final dark relaxation.

### JTS script for determining ECS

5(4(100msD)100ms/X=A[on]10usA[off]160usB[on]10usB[off]/2(300usD)400usD1msD3msD3(5ms D)4(20msD)4(50msD)2(100msD)4(500msD)3(1sD)0(5sD))2usH2(10msD)T2usH2500ms2usG[7673 4uA]2500ms2usH2500msG[76734uA]6ms/Z=5(750usH140usD2usG[76734uA])2usH1948usD2usH4 (750usH140usD2usH)2usG[76734uA]2us/31msZ41msZ191msZ196msZ191msZ191msZ290msZ390 msZ390msZ2usH2500ms5(750usH140usD2usH)2usG[76734uA]5(750usH140usD2usG[76734uA])2 1msZ41msZ116msZ191msZ191msZ191msZ191msZ384msZ408msZ692msZ2usH2500ms5(2usG[76 734uA]6msZ31msZ41msZ191msZ196msZ191msZ191msZ290msZ390msZ390msZ2usH2500ms5(75 0usH140usD2usH)2usG[76734uA]5(750usH140usD2usG[76734uA])21msZ41msZ116msZ191msZ1 91msZ191msZ191msZ384msZ408msZ692msZ2usH2500ms)

**Supplemental Table S2.**
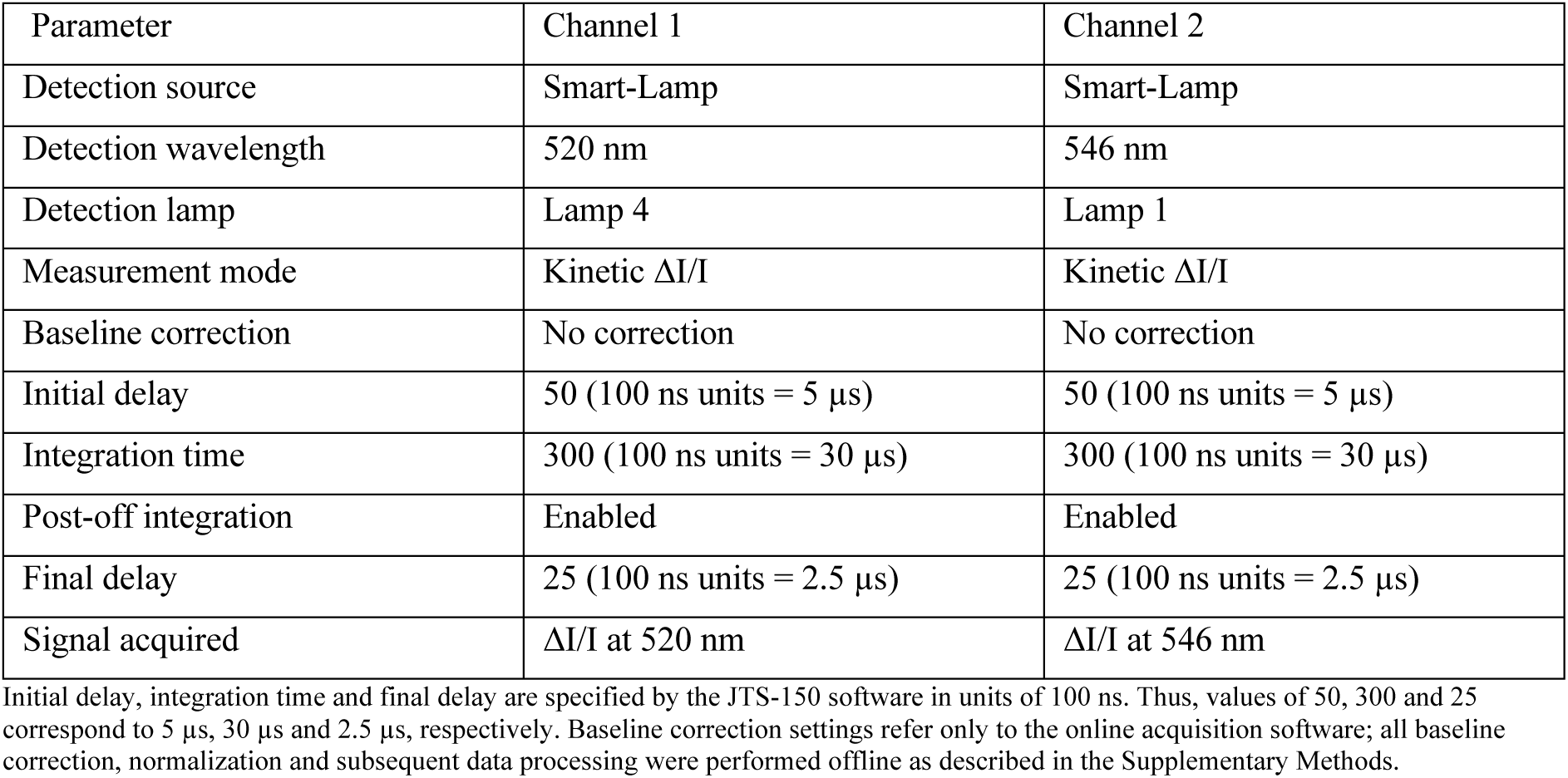
JTS-150 Detection Settings Used for ECS Measurements.

| Parameter | Channel 1 | Channel 2 |
| --- | --- | --- |
| Detection source | Smart-Lamp | Smart-Lamp |
| Detection wavelength | 520 nm | 546 nm |
| Detection lamp | Lamp 4 | Lamp 1 |
| Measurement mode | Kinetic $\Delta I/I$ | Kinetic $\Delta I/I$ |
| Baseline correction | No correction | No correction |
| Initial delay | 50 (100 ns units = 5 μs) | 50 (100 ns units = 5 μs) |
| Integration time | 300 (100 ns units = 30 μs) | 300 (100 ns units = 30 μs) |
| Post-off integration | Enabled | Enabled |
| Final delay | 25 (100 ns units = 2.5 μs) | 25 (100 ns units = 2.5 μs) |
| Signal acquired | $\Delta I/I$ at 520 nm | $\Delta I/I$ at 546 nm |

Initial delay, integration time and final delay are specified by the JTS-150 software in units of 100 ns. Thus, values of 50, 300 and 25 correspond to 5 μs, 30 μs and 2.5 μs, respectively. Baseline correction settings refer only to the online acquisition software; all baseline correction, normalization and subsequent data processing were performed offline as described in the Supplementary Methods.

Detection settings. ECS measurements were performed using the Smart-Lamp detection module of the JTS-150. Absorbance changes were monitored simultaneously at 520 and 546 nm in Kinetic ΔI/I mode. Both detection channels were configured with an initial delay of 50 (5 µs), an integration time of 300 (30 µs), and a final delay of 25 (2.5 µs). Baseline correction was disabled within the acquisition software, whereas post-off integration was enabled for both channels. All baseline correction and normalization were subsequently performed during offline data analysis.

### Single-turnover ECS measurements

Prior to DIRK measurements, ECS kinetics induced by a single-turnover flash were recorded. Laser firing was controlled through a two-stage trigger sequence consisting of a flashlamp trigger followed by a 160 µs delay and a subsequent Q-switch trigger. The resulting nanosecond laser pulse-initiated charge separation in photosystems.

Immediately after excitation, ECS relaxation kinetics were sampled using progressively increasing acquisition intervals. Measurements were collected at approximately 0.3, 0.6, 1, 2, 5, 10, 15, 20, 40, 60, 80, 100, 150, 200, 250, 300, 400, 500, 1000, 1500, 2000, 2500, 3500, 4500 and 5500 ms after the laser flash. The complete single-turnover acquisition sequence was repeated five times and averaged.

### Light adaptation and establishment of steady-state electron transport

Following single-turnover measurements, samples were exposed to continuous actinic illumination for 2.5 s using the actinic LED source. This illumination period was used to establish steady-state photosynthetic electron transport and generate a stable proton motive force. Actinic illumination was subsequently interrupted, and samples were allowed to relax in darkness for an additional 2.5 s. The acquisition time axis was then reset, and all subsequent DIRK measurements were referenced relative to this new time origin.

### DIRK measurements

Dark-interval relaxation kinetics (DIRK) were acquired by periodically interrupting continuous actinic illumination with short dark intervals. Each DIRK acquisition consisted of: 750 µs continuous actinic illumination, followed by a 140 µs dark interruption, during which ECS absorbance was measured, after which actinic illumination was restored.

Five consecutive DIRK acquisitions were performed under illumination. An additional set of five DIRK acquisitions was subsequently performed during the following dark interval. Together these ten measurements constituted one Z-frame acquisition block.

### Z-frame acquisition sequence

Following establishment of steady-state illumination, an initial Z-frame was acquired immediately at *t=0*.

Additional Z-frame acquisitions were subsequently performed following delays of 31, 41, 191, 196, 191, 191, 290, 390 and 390 ms.

After a 2.5 s dark recovery period, five dark DIRK acquisitions followed by five illuminated DIRK acquisitions were recorded. A second series of Z-frame acquisitions was then performed following delays of 21, 41, 116, 191, 191, 191, 191, 384, 408 and 692 ms.

This complete acquisition cycle was repeated five times. The total duration of the ECS experiment was approximately 95–100 s.

### ECS data processing

Raw ECS traces exported from the JTS-150 were first converted from seconds to milliseconds. The time axis was then shifted such that the first laser flash occurred at *t=0*. Because the acquisition script contains approximately 400 ms of dark measurements followed by an additional 100 ms delay before the first flash, all traces were aligned relative to the first flash event. For each kinetic point, the five technical repetitions generated by the acquisition script were averaged. The averaged signal was subsequently normalized to the mean ECS signal recorded during the dark baseline immediately preceding laser excitation.

The ECS amplitude generated under DCMU/HA inhibition was used as the PSI-only reference signal. The ratio between the ECS signal recorded under experimental conditions and the ECS signal measured under DCMU/HA conditions was used to estimate the relative contribution of PSII versus PSI charge separation as a function of time after the laser flash.

### DIRK analysis

For DIRK measurements, the original time axis was converted to milliseconds and shifted by 7.5 s, corresponding to the adaptation phase preceding initiation of the DIRK sequence. Following this transformation, the first DIRK acquisition was defined as *t = 0*.

For each acquisition point, values from all technical repetitions were averaged. The averaged traces were subsequently corrected by subtracting the mean dark baseline measured immediately before the laser-flash sequence.

Dark-interval relaxation rates were determined from the initial ECS decay during each DIRK event. For every DIRK measurement, a linear least-squares regression was calculated using the first five acquisition points collected during the corresponding dark interval. The regression was performed using the actual acquisition times of the first five data points within each DIRK event. Electron transfer rates were calculated as the difference between the slope measured under illumination and the slope measured during the corresponding dark interval:

Electron transfer rate = slope_light_ – slope_Dark_

The resulting value was normalized to the maximal PSI-only ECS signal measured after the laser flash in DCMU/HA-treated samples.

Final values were expressed as electrons per second per PSI [(*e*^−^) ⋅ *s*^−1^ ⋅ *PSI*^−1^] corresponding to the number of electrons transferred per second per PSI reaction center. Although the complete JTS acquisition sequence is shown for reproducibility, only the single-turnover flash kinetics and the initial D2 DIRK relaxation slope were used for the analyses presented in the manuscript. The remaining acquisition points are included solely to document the complete measurement protocol.

**Supplemental Figure S7.**
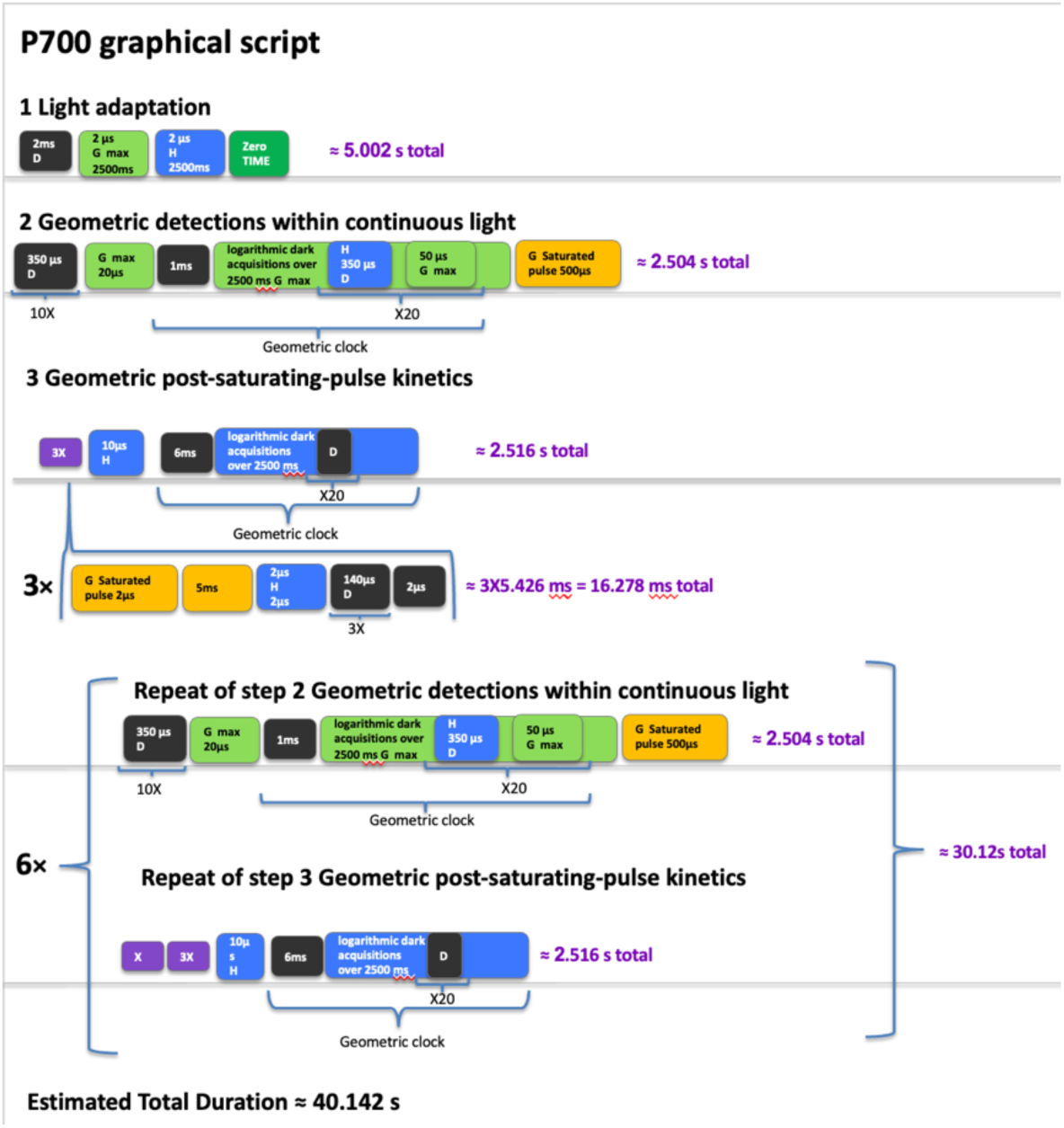
Graphical representation of the JTS script for Photosystem I (P700) redox kinetics and quantum yield measurements. (1) Pre-Illumination Baseline: Measures the dark-adapted sample to establish the initial baseline before any light treatment begins. (2) Actinic Induction and Steady State: Exposes the cells to continuous actinic illumination to record the steady-state P700 oxidation level (Ps). (3) Saturating Pulse Phase: Delivers a brief, intensive multi-turnover light flash to transiently drive the remaining functional P700 pool to its maximum oxidation level (Pm). (4) Dark Recovery Phase: Tracks the final dark reduction kinetics under detection light only to capture the post-pulse recovery profile.

### JTS script for determining P700

2msD1(2usG[42000uA]2500ms2usH2500ms)T10(350usD)G[42000uA]20us

{1ms,20,2500ms,H350usD50usG[42000uA]}G[1000000uA]500us

/X=G[1000000uA]2us5ms2usH2us3(140usD)2us/ 3(X)10usH{6ms,20,2500ms,D}6(10(350usD)G[42000uA]20us

{1ms,20,2500ms,H350usD50usG[42000uA]}G[1000000uA]500us X3(X)10usH{6ms,20,2500ms,D})

**Supplemental Table S3.** JTS-150 Detection Settings Used for P700 Measurements.

| Parameter | Channel 1 | Channel 2 |
| --- | --- | --- |
| Detection source | Smart-Lamp | Smart-Lamp |
| Detection wavelength | 705 nm | 740 nm |
| Detection lamp | Lamp 7 | Lamp 2 |
| Measurement mode | Kinetic $\Delta I/I$ | Kinetic $\Delta I/I$ |
| Baseline correction | No correction | No correction |
| Initial delay | 50 (100 ns units = 5 μs) | 50 (100 ns units = 5 μs) |
| Integration time | 300 (100 ns units = 30 μs) | 300 (100 ns units = 30 μs) |
| Post-off integration | Disabled | Disabled |
| Final delay | 25 (100 ns units = 2.5 μs) | 25 (100 ns units = 2.5 μs) |
| Signal acquired | $\Delta I/I$ at 705 nm | $\Delta I/I$ at 740 nm |

Final analytical signal: $\Delta I/I_{705-740}$ calculated from the two detection channels.

### Detailed P700 acquisition sequence

P700 kinetics were acquired using a custom JTS-150 timing script. A graphical representation of the acquisition sequence is provided in Supplemental Figure S7, and the complete instrument script below it.

### Light adaptation and dark recovery

Prior to kinetic measurements, samples were exposed to continuous actinic illumination (42,000 µA LED driver setting) for 2.5 s. Illumination was then interrupted and samples were maintained in darkness for an additional 2.5 s. Following this adaptation procedure, the acquisition time axis was reset, and all subsequent measurements were referenced relative to this new time origin.

### Baseline acquisition

Immediately following time-axis reset, ten absorbance measurements were acquired at intervals of 350 µs. These measurements were used for baseline determination and signal correction.

### Geometric acquisition under illumination

Continuous actinic illumination was restored and maintained throughout the first acquisition phase. P700 kinetics were recorded using a geometric acquisition schedule defined as:

{1 ms, 20 acquisitions, 2500 ms total duration}

Twenty measurements were collected over a total period of 2.5 s with logarithmically increasing intervals between acquisitions. Each acquisition event consisted of: interruption of actinic illumination, a 350 µs dark interval, absorbance acquisition during the dark interval, restoration of actinic illumination 50 µs later.

This acquisition mode allowed monitoring of P700 redox state while maintaining continuous photosynthetic activity.

### Saturating pulse

At the end of the illumination phase, a saturating pulse was delivered using the actinic LED source operated at a driver setting of 1,000,000 µA. Pulse duration was 500 µs.

### Post-saturating probing sequence

Immediately after the saturating pulse, a probing sequence (X-frame) was executed three consecutive times. Each X-frame consisted of: saturating illumination, 5 ms illumination period, interruption of illumination, three consecutive 140 µs dark intervals, absorbance measurements during the dark intervals.

These measurements were used to monitor rapid P700 reduction kinetics immediately following maximal oxidation.

### Geometric recovery kinetics

Following completion of the X-frame sequence, P700 reduction kinetics were monitored using a second geometric acquisition schedule: {6 ms, 20 acquisitions, 2500 ms total duration} Twenty measurements were collected over a total period of 2.5 s using logarithmically increasing intervals. This second geometric schedule monitored the recovery of the P700 redox state after the saturating pulse.

### Repeated acquisition cycles

After completion of the initial acquisition cycle, the following sequence:

Baseline acquisition, geometric measurements during illumination, saturating pulse, X-frame probing sequence, geometric recovery kinetics, was repeated six additional times. Consequently, a total of seven acquisition cycles were recorded for each sample.

### Data processing

P700 traces were first baseline corrected using the average of the ten pre-illumination measurements acquired immediately after time reset, averaged across technical repetitions, and normalized to the maximal P700 oxidation measured under DCMU/HA inhibition.

The maximal oxidizable PSI pool (P′m) was defined as the highest normalized P700 signal observed during the saturating-pulse region in DCMU/HA-treated samples.

The maximal P700 oxidation under experimental conditions (Pm) was defined as the highest normalized P700 signal observed during the saturating-pulse region of the corresponding untreated trace.

Steady-state P700 oxidation (Ps) was defined as the P700 level immediately preceding application of the saturating pulse under continuous actinic illumination.

The effective PSI quantum yield (YI), donor-side limitation (YND), and acceptor-side limitation (YNA) were calculated according to:

YI = (Pm − Ps) / P′m

YND = Ps / P′m

YNA = (P′m − Pm) / P′m

All calculations were performed using traces averaged across technical repetitions.

**Supplemental Table S4.** Data processing parameters used for ECS, DIRK and P700 analyses.

| Analysis step | Parameter used |
| --- | --- |
| ECS time conversion | Time converted from seconds to milliseconds |
| ECS alignment | First laser flash defined as $t = 0$ |
| ECS baseline | Average of all dark points preceding first laser flash |
| ECS averaging | Five technical repetitions averaged for each time point |
| ECS normalization | Divided by PSI-only ECS amplitude measured in DCMU/HA-treated samples |
| PSI/PSII ratio | Calculated from total ECS amplitude minus PSI-only ECS amplitude |
| DIRK time conversion | Time converted to milliseconds |
| DIRK alignment | 7.5 s adaptation period subtracted; first DIRK acquisition defined as $t = 0$ |
| DIRK averaging | Corresponding points averaged across all technical repetitions |
| DIRK baseline correction | Average dark signal immediately preceding each DIRK block subtracted |
| DIRK slope determination | Initial linear slope extracted from each DIRK relaxation trace |
| Electron transfer rate | Light slope – dark slope |
| PSI normalization | Electron transfer rate divided by maximal PSI signal measured under DCMU/HA conditions |
| Final DIRK units | electrons $s^{-1}$ PSI$^{-1}$ |
| P700 baseline (P0) | Signal measured in darkness before illumination |
| Ps determination | Steady-state P700 level immediately before saturating pulse |
| Pm determination | Maximal P700 oxidation during saturating pulse |
| P'm determination | Maximal oxidizable P700 pool in presence of DCMU + HA |
| YI calculation | $(P_m - P_s)/P'm$ |
| YNA calculation | $(P'm - P_m)/P'm$ |
| YND calculation | $P_s/P'm$ |
| CEF calculation | $k_{ox} \times [P700_{red}]$ |
| [P700red] calculation | $(P_m - P_s)/P'm$ |
| kox determination | Derived independently from ECS measurements |

**Supplemental Table S5.** Parameter extraction rules.

| Parameter | Definition |
| --- | --- |
| Ps | Signal at the last acquisition point before the saturating pulse (steady state under actinic light) |
| Pm | Maximum signal during saturating pulse |
| P'm | Maximum signal under DCMU/HA |
| ECS PSI-only amplitude | First post-flash point under DCMU/HA |
| DIRK slope | Linear fit of first five points after dark interruption |
| ETR | Light slope – dark slope |
| CEF | $k_{ox} \times [P700red]$ |

### MIMS data analysis

Raw ion currents for hydrogen (H_2_), oxygen (O_2_), and carbon dioxide (CO_2_) were extracted from the MIMS datasets. To convert the raw currents into absolute chemical concentrations, linear calibration curves were generated immediately adjacent to the experimental measurements using known gas standards.

### Hydrogen Calibration

Standard curves were constructed using 100% H_2_ gas generated from a hydrogen generator, which was subsequently diluted by half (50% H_2_) for the high-concentration injections, alongside a 5% H_2_ gas standard to define the lower boundaries of the linear range.

### Oxygen Calibration

Ambient air was used to establish the 21% O_2_ saturation point, while a zero-oxygen reference (0% O_2_) was obtained by thoroughly purging the system with an inert gas bottle.

The slope and intercept parameters derived from these standard curves were used to calculate relative gas abundances, which were subsequently converted to micromolar (μM) gas concentrations and normalized to the total chlorophyll concentration of the sample. Kinetic traces were baseline-corrected to set both the time and the gas concentration to zero at the exact onset of the light induction period (*t=0*). Final data profiles represent the mean values calculated from at least three independent biological replicates, with variance expressed as standard error of the mean (SEM).

## Notes

### Competing Interest Statement

The authors have declared no competing interest.

